# Damage activates *EVG1* to suppress vascular differentiation during regeneration in *Arabidopsis thaliana*

**DOI:** 10.1101/2023.02.27.530175

**Authors:** Shamik Mazumdar, Ai Zhang, Constance Musseau, Muhammad Shahzad Anjam, Peter Marhavy, Charles W. Melnyk

## Abstract

Plants possess remarkable regenerative abilities to form *de novo* vasculature after damage and in response to pathogens that invade and withdraw nutrients. To look for common factors that affect vascular formation upon stress, we searched for *Arabidopsis thaliana* genes differentially expressed during *Agrobacterium* infection, nematode infection and plant grafting. One such gene was cell-wall associated and highly induced by all three stresses. Mutations in it enhanced ectopic xylem formation in Vascular cell Induction culture System Using Arabidopsis Leaves (VISUAL) and enhanced graft formation and was thus named *ENHANCER OF VISUAL AND GRAFTING 1* (*EVG1*). Mutated *evg1* inhibited cambium development and callus formation yet promoted tissue attachment, syncytium size, phloem reconnection and xylem formation. *evg1* affected abscisic acid and cell wall responses and was itself down regulated by ABA. We found mutations in a receptor-like gene, *RLP44*, had the same regeneration phenotype as *EVG1* mutations including enhancing VISUAL and grafting. *evg1* and *rlp44* mutants affected the expression of many genes in common including those important for successful regeneration and vascular formation. We propose that *EVG1* integrates information from cutting, wounding or parasitism stresses and functions with *RLP44* to suppress vascular differentiation during regeneration.

## Introduction

The ability for plants to regenerate tissues after injury is of fundamental importance to maintain tissue integrity and regrow. Upon wounding, plants activate defence and regeneration responses to deter further injury and to heal damage. Wounding induces cell wall damage, causes auxin accumulation and increases auxin response around the injury, processes thought to be critical for downstream activation of transcriptional factors *ERF115, TMO6* and *HCA2* important for wound healing (Canher et al., 2020; Hoermayer et al., 2020; Zhang et al., 2022). During early stages of regeneration, cells close to the site of wounding expand and deposit cell wall material such as pectins which can help tissues adhere (Sala et al., 2019; Zhang et al., 2022). Cells dedifferentiate and divide to form a mass of pluripotential stems cells known as callus which is regulated by cell cycle genes including *CYCLIN D3;1* (*CYCD3;1*) and the transcription factor *WOUND INDUCED DIFFERENTIATION 1* (*WIND1*) that activates cytokinin responses (Iwase et al., 2011a; Ikeuchi et al., 2017a). Callus tissues fill the wound and differentiate to reform missing cell types and vascular connections by differentiating new phloem and xylem (Iwase et al., 2021).

Similar recognition and healing processes occur during the horticulturally relevant process of plant grafting when two plants are cut and joined together. There, thousands of genes are differentially expressed at the graft junction with cambium-related genes activating first followed by phloem-related and lastly xylem-related genes (Melnyk et al., 2018). Several genes have been identified as important for graft formation, namely the auxin-related genes *ALF4* and *AXR1* that are needed below the graft junction, and cambium-related genes such as *HCA2, TMO6, WOX4* and *ANAC096* (Melnyk et al., 2015, 2018; Zhang et al., 2022; Thomas et al., 2022; Matsuoka et al., 2016). Mutating these factors reduces vascular connectivity or cambium formation, but to date, no recessive mutations that improve grafting have been identified.

Given the importance of cell walls during regeneration, it is surprising that no role has been found yet for brassinosteroids (Nanda and Melnyk, 2018), a group of plant hormones involved in vascular development and cell wall homeostasis (Caño-Delgado et al., 2004; Ibañes et al., 2009; Wolf et al., 2012; Lozano-Elena and Caño-Delgado, 2019; Oh et al., 2020). Activating brassinosteroid signaling is critical for forming ectopic xylem from mesophyll cells in leaves through the Vascular Cell Induction Culture System Using Arabidopsis Leaves (VISUAL) system (Kondo et al., 2015). By enhancing or suppressing brassinosteroid signaling, more or less ectopic xylem is formed (Kondo et al., 2015). However, during normal root development, mutations in the brassinosteroid receptor BRASSINOSTEROID INSENSITIVE 1 (BRI1) promote xylem formation and suppress cambium, yet this role seems independent of canonical brassinosteroid signaling and instead is related to the interaction of BRI1 with RECEPTOR-LIKE PROTEIN 44 (RLP44). There, BRI1-RLP44 associates with the phytosulfokine pathway to promote cambial divisions but repress xylem formation (Holzwart et al., 2018). Phytosulfokine signaling is known to be activated by wound-induced *ERF115* (Heyman et al., 2013) yet the role of canonical and non-canonical brassinosteroid signaling during grafting and regeneration remains poorly characterized.

Although defence responses are induced by most pathogens, some plant pathogens also activate regeneration pathways to infect their hosts more efficiently. *Agrobacterium* enters plant wound sites and incudes auxin and cytokinin production to cause cell differentiation, cell division, vascularization and tumour growth (Deeken et al., 2007; Zhang et al., 2015). Root-knot nematodes and cyst nematodes feed on plant roots and cause proliferation of cells to derive nutrients from the host plants (Shanks et al., 2016; Olmo et al., 2020). They also activate host genes that regulate vascular development such as *HOMEOBOX-8* (*ATHB8*), *WUSCHEL-RELATED HOMEOBOX 4* (*WOX4*), and *TRACHEARY ELEMENT DIFFERENTIATION INHBITORY FACTOR RECEPTOR* (*TDR/PXY*) (Yamaguchi et al., 2017). The cyst nematode *Heterodera schachtii* releases CLE-like effector proteins into plant cells which induces cell proliferation and activates the *WOX4* cambium-promoting pathway (Guo et al., 2017). Thus, similarities such as *WOX4* activation during nematode infection and grafting suggests an overlap in common processes induced during various forms of regeneration or parasitism (Melnyk, 2017a). However, what genes regulate these common processes and how they promote or inhibit parasitism and regeneration remains largely unknown. Here, we further investigate these aspects to identify and characterize a gene, AT3G08030, previously called *ATHA2-1* due to phylogenetic relatedness to a clade of DUF642 protein genes (Vázquez-Lobo et al., 2012a). AT3G08030 mutants enhance vascular formation during both grafting and VISUAL. Mutating this gene affects multiple regeneration and developmental aspects and appears similar to mutations in RLP44 in regulating development and regeneration. Given that AT3G08030 is activated highly upon wounding or parasitism, we propose it acts as a stress responsive gene that balances cambium and *de novo* vascular formation.

## Results

### *EVG1* responds to stress and regulates regeneration

To identify genes differentially expressed in response to stress, we compared previously published *Arabidopsis thaliana* transcriptomic datasets from *Agrobacterium* infection, nematode infection and plant grafting (Deeken et al., 2007; Szakasits et al., 2009; Barcala et al., 2010; Melnyk et al., 2018). Of those differentially expressed, we selected for genes associated with development and narrowed our list to 22 candidates which included previously described vascular-related genes such as *ATHB8* (AT4G32880) and *WRKY23* (AT2G47260) (Baima et al., 2001; Prát et al., 2018) (Figure 1A) As a secondary screening method to find novel vascular regulators, we employed the VISUAL system of ectopic xylem formation that could rapidly identify mutants associated with vascular development (Kondo et al., 2015). T-DNA mutant lines were tested using VISUAL and many mutant genes reduced ectopic xylem formation consistent with a role for these genes in promoting vascular development (Figure 1B). However, AT3G08030 appeared exceptional since mutating this gene increased levels of ectopic xylem formation (Figure 1B). This gene was highly upregulated early during graft formation (Figure 1C) and previously published datasets showed that AT3G08030 was upregulated by abiotic stresses including osmotic, salt, heat, drought, cold, and UV-B, and biotic stress such as *Heterodera schachtii* infection (Supplemental Figure 1, A and B) (data from ePlant Browser (Bar Toronto) (Kilian et al., 2007; Fucile et al., 2011; Waese et al., 2017)

**Figure 1.**
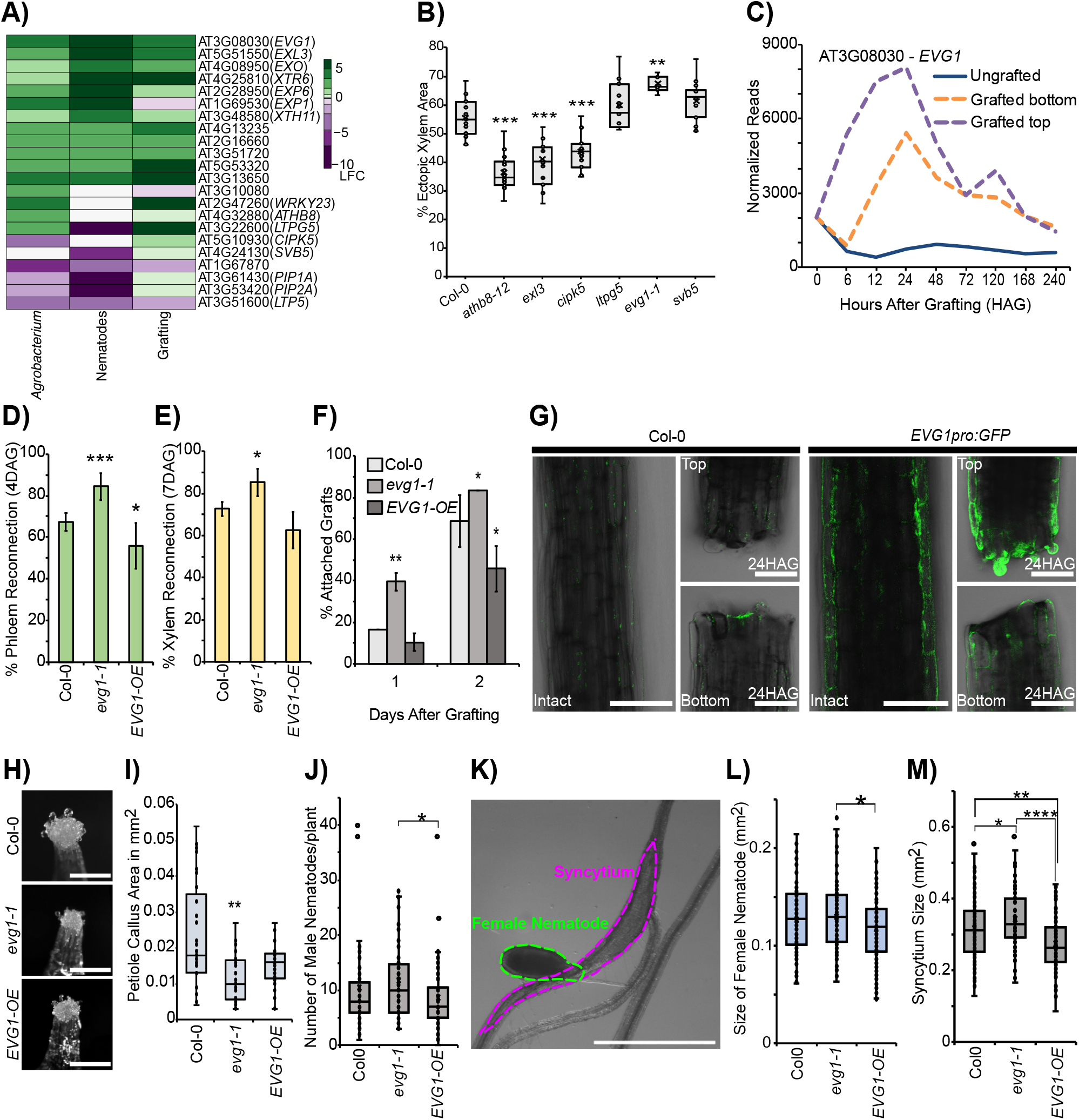
*EVG1* responds to damage and regulates grafting and callus formation. (A) Heat map showing differentially expressed genes in *Agrobacterium* infected host (Deeken et al., 2007), nematode infected host (Szakasits et al., 2009; Barcala et al., 2010) and grafted top vs ungrafted at 24 hours (Melnyk et al., 2018). (B) VISUAL assay quantifications of ectopic xylem area in WT and six loss of function mutants of interest. Dots represent individual samples (n=15). Asterisks indicate significant differences compared to Col-0. ** p<0.001, *** p<0.001, Wilcoxon’s test. (C) Dynamics of *EVG1* expression during grafting (Melnyk et al., 2018). (D-E) Reconnection percentage of phloem (4 days after grafting – DAG) and xylem (7 DAG) in Col-0, *evg1-1* and *EVG1-OE*. The mean ± SD of 5 experiments with n=12 per genotype per experiment is shown. Asterisks indicate significant differences compared to Col-0. * p<0.05, ** p<0.01, Student’s t-test. (F) Percentage of attached grafts of Col-0, *evg1-1* and *EVG1-OE* at 1 DAG and 2 DAG. The mean ± SD of 4 experiments with n=12 per genotype per experiment is shown. Asterisks indicate significant differences compared to Col-0. * p<0.05, ** p<0.01, Student’s t-test. (G) Comparison of intact and grafted tissues of Col-0 and *EVG1pro:GFP* 24 hours of grafting. Scale bars, 100 μm. (H) Representative images of callus formation in petiole explants of Col-0, *evg1-1* and *EVG1-OE*. Scale bar, 250 μm. (I) Petiole callus areas of Col-0, *evg1-1* and *EVG1-OE*. Dots represent individual samples (n=29 per genotype). Asterisks show statistical significance compared to Col-0. ** p<0.01, Wilcoxon’s test. (J) Average number of male nematodes per plant 12 days post infection (dpi) in Col-0, *evg1-1* and *EVG1-OE*. Dots indicate individual samples. * p<0.05, Wilcoxon’s test. (K) Picture showing infection of Col-0 root by a female *Heterodera schachtii* nematode. Green dotted line indicates female size, magenta dotted line indicates syncytium size. Scale bar represents 1mm. (L-M) Average size of female nematodes per plant and average syncytium size at 14 dpi in Col-0, *evg1-1* and *EVG1-OE*. Dots indicate individual samples. * p<0.05, ** p<0.01, **** p<0.001, Wilcoxon’s test.

Since AT3G08030 was highly upregulated during graft formation (Figure 1C), we tested whether this gene affects *Arabidopsis* hypocotyl graft formation dynamics and vascular regeneration. We applied the vascular mobile dye carboxyfluorescein diacetate (CFDA) to grafted scions or rootstocks using previously described assays (Melnyk et al., 2015; Melnyk, 2017c). We found that the mutant showed enhanced phloem and xylem reconnection compared to wild type Col-0 plants (Figure 1, D and E). Since this mutant enhanced both VISUAL and grafting, we named AT3G08030 as *ENHANCER OF VISUAL AND GRAFTING* (*EVG1*) and the mutation as *evg1-1*. To further test the role *EVG1* in grafting, we employed a gain of function strategy and obtained an over expressing *35Spro:EVG1-cDNA* line (*EVG1-OE*) from the FOX hunting system (Ichikawa et al., 2006). *EVG1-OE* significantly reduced phloem connectivity (Figure 1D, Supplemental Figure 1C) but had an insignificant effect on xylem connectivity when compared to Col-0 (Figure 1E, Supplemental Figure 1D). We checked tissue attachment during grafting since attachment is needed for phloem and xylem connection (Melnyk et al., 2015; Melnyk and Meyerowitz, 2015). *evg1-1* improved attachment at the graft junction whereas *EVG1-OE* reduced tissue attachment compared to Col-0 (Figure 1F). To identify where at the junction *EVG1* played a role, we performed heterografting assays and found that *evg1-1* grafted as a scion on a wild type Col-0 rootstock enhanced grafting whereas the reciprocal combination reduced grafting efficiency. Overexpressing *EVG1* either in the scion or the rootstock reduced grafting efficiency (Supplemental Figure 1E). We generated a *EVG1pro:GFP* transcriptional reporter and found it activated at the graft junction after cutting, particularly in the scion (Figure 1G), consistent with previous transcriptome sequencing at the junction (Figure 1C) (Melnyk et al., 2018). However, *EVG1* activation was not specific to grafting and also occurred after hypocotyl wounding with forceps (Supplemental Figure 1G). Since graft formation typically involves callus formation (Melnyk et al., 2015; Melnyk and Meyerowitz, 2015), we tested whether callus formation required *EVG1. evg1-1* reduced wound-induced callus formation from petioles when compared to wild type Col-0, however *EVG1-OE* had little effect (Figure 1, H and I). In hypocotyl callus, *evg1-1* had an insignificant effect (Supplemental Figure 1F).

To test the role of *EVG1* beyond wounding or grafting, we performed infection assays with the plant-parasitic cyst nematode, *Heterodera schachtii*. Nematodes characteristically develop feeding sites near the vasculature and induce *de novo* phloem formation, assumed to symplastically connect the syncytial feeding structures to the vascular bundles for continuous supply of nutrients (Melnyk et al., 2017a). 12 days post infection, the total number of male nematodes were significantly increased in *evg1-1* compared to *EVG1-OE* (Figure 1J). Female nematodes did not change in number but were reduced in size in *EVG1-OE* (Figure 1, K and L; Supplemental Figure 1, H and I). At the infection site, *evg1-1* had larger syncytium than wild type Col-0 whereas in *EVG1-OE*, the syncytium size was reduced compared to wild type Col-0 (Figure 1M). Taken together, our results show that *EVG1* was activated by stress and played a role in suppressing graft formation and syncytium development.

### *EVG1* expresses in epidermal cell layers and regulates vascular development

To understand the role of *EVG1* during non-stressed conditions, we analysed transcriptional and translational *EVG1* fluorescent reporters. In *Arabidopsis* primary roots, fluorescence was highest in the root epidermis with little signal in the cortex or stele (Figure 2, A and B, Supplemental Figure 2, A and B). Previously published datasets confirmed the expression of *EVG1* in outer cell layers of the root (Supplemental Figure 2C) (https://rootcellatlas.org/) (Ryu et al., 2019; Wendrich et al., 2020; Denyer et al., 2019; Jean-Baptiste et al., 2019; Zhang et al., 2019; Shulse et al., 2019; Shahan et al., 2022). However, in lateral root primordia, we observed strong *EVG1* expression throughout inner and outer cell layers (Figure 2C). The subcellular localization of *EVG1pro:EVG1-GFP* suggested that EVG1 may be present in the plasma membrane or extracellular regions of the cell (Figure 2D) consistent with protein predictions (Figure 2E) (Thumuluri et al., 2022). To investigate the role of *EVG1* in vascular development, we obtained a second *EVG1* mutant allele, *evg1-2*, that showed even lower *EVG1* transcript levels than *evg1-1* (Supplemental Figure 2, D and E). In VISUAL assays for xylem formation, *evg1-2* enhanced ectopic xylem formation like *evg1-1* whereas *EVG1-OE* reduced ectopic xylem formation compared to WT (Figure 2, F and G). During VISUAL, (pro)cambium related genes peak at 24 hours post induction and subsequently reduce at later time points (Kondo et al., 2015). We found that *EVG1* transcript levels were also elevated at 24-hours and reduced as time progressed (Figure 2H). VISUAL induces ectopic xylem formation in part by enhancing brassinosteroid signaling (Kondo et al., 2014, 2015). We tested how brassinosteroid application affected *evg1-1*. Xylem identity in *evg1-1* primary roots appeared normal but when treated with epiBrassinolide (epiBL), *evg1-1* had reduced extra protoxylem in outer metaxylem and reduced reticulate metaxylem when compared to treated wild-type Col-0 (Figure 2I). We also check the number of metaxylem cell files in *evg1-1* primary roots and found additional metaxylem cell files in *evg1-1* compared to wild type (Figure 2J). Since *EVG1* showed a similar expression pattern during grafting and VISUAL as genes associated with cambium formation, we checked cambium development in *evg1* (Kondo et al., 2015; Melnyk et al., 2018). Cross-sections below the hypocotyl-root junction showed reduced cambium and xylem area in *evg1-1* compared to wild-type Col-0 (Figure 2, K-M). We did not observe changes in the cambium to xylem area ratio between Col-0 and *evg1-1*, but when normalized for unit xylem area, *evg1-1* had more xylem cells per unit area than Col-0 (Supplemental Figure 2, G-J). Taken together, these results suggest that *EVG1* promoted cambium formation but repressed xylem differentiation and cell expansion.

**Figure 2.**
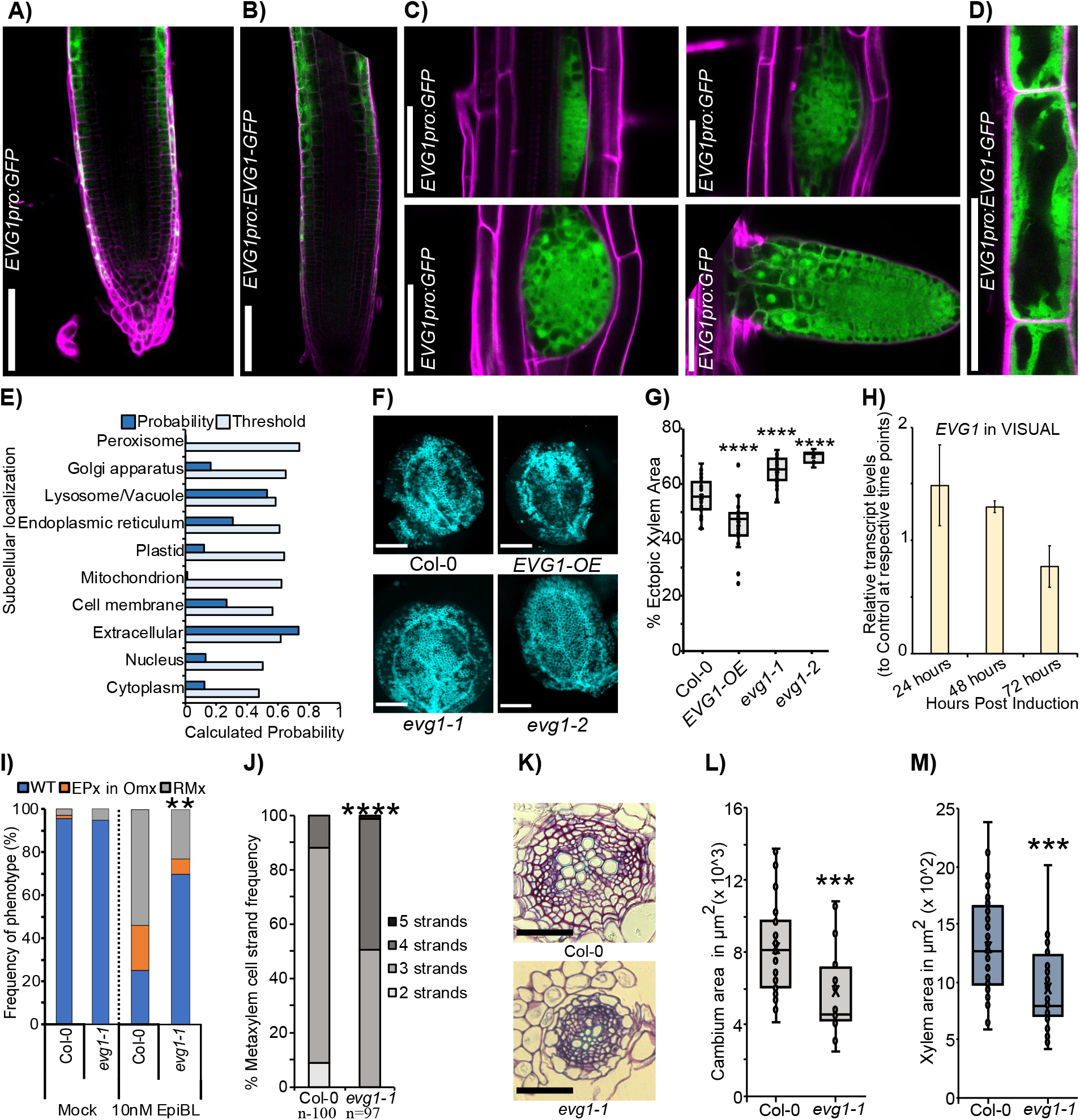
*EVG1* regulates vascular development. (A-B) *EVG1pro:GFP* and *EVG1pro:EVG1-GFP* fluorescence at the root tip. Cell walls stained by PI (magenta). Scale bar represents 100 μm. (C) *EVG1pro:GFP* fluorescence in different stages of lateral root development. Cell walls stained by PI (magenta). Scale bar represents 100 μm. (D) *EVG1pro:EVG1-GFP* fluorescence in root tip epidermal cells. Cell walls stained by PI (magenta). Scale bar represents 50 μm. (E) Bar plot showing the threshold and calculated probability of subcellular localization of the EVG1 protein, predicted using DeepLoc2.0 (Thumuluri et al., 2022). (F) VISUAL assay images of ectopic xylem formation in Col-0, *evg1-1, evg1-2* and *EVG1-OE*, scale bars 1 mm. (G) Ectopic xylem quantifications of Col-0, *evg1-1* and *EVG1-OE*. Dots represent individuals Col-0 (n=51), *evg1-1* (n=46), *evg1-2* (n=10), *EVG1-OE* (n=24). Asterisk show statistical significance compared to Col-0, ****, p<0.0001, Wilcoxon’s test. (H) Transcript levels of *EVG1* during VISUAL induction. (I) Xylem phenotypes of Col-0 (n=50) and *evg1-1* (n=50) under mock and exogenous epiBL treatment: wild type (WT), extra protoxylem in outer metaxylem (EPx in Omx) or reticulate metaxylem (RMx). Asterisks indicate significant difference compared to epiBL treated Col-0. **, p<0.001, ****, p<0.0001, Fisher’s exact test with Benjamini-Hochberg adjustment. (J) Metaxylem strand number in Col-0 and *evg1-1*. Asterisk show statistical significance compared to Col-0, ****, p<0.0001, Fisher’s exact test with Benjamini-Hochberg adjustment. (K) Cross sections 2 mm below 21-day old shoot-root junction of Col-0 and *evg1-1* seedlings. Scale bars, 200 μm. (L-M) Cambium area and xylem area quantifications of Col-0 and *evg1-1*. Dots represent individual Col-0 (n=35) and *evg1-1* (n=24). Asterisk show significant difference compared to Col-0. *** p<0.001, Wilcoxon’s test.

### *RLP44* mutants phenocopy *EVG1* mutants during regeneration

Our finding that *evg1-1* induced more xylem in VISUAL prompted us to investigate the role of brassinosteroids more closely during regeneration. We tested the effects of mutations in brassinosteroid signaling related genes *BRASSINOSTEROID INSENSITIVE 1* (*BRI1*), *BRASSINOSTEROID INSENSITIVE 2* (*BIN2*), *BRI1-EMS SUPPRESOR 1* (*BES1*), *BRASSINAZOLE RESISTANT 1* (*BZR1*) and *RLP44* in grafting and callus formation assays. Grafting *bri1-301, bes1-2, bzr1-D* and *bin2-1* reduced phloem and xylem reconnection but exceptionally, *rlp44-3* increased rates of phloem connectivity compared to wild type Col-0 and had a slight but non-significant increase in xylem connectivity (Figure 3, A and B; Supplemental Figure 3A). We obtained a *35Spro:RLP44-RFP (RLP44ox)* line (Wolf et al., 2014) and found it behaved oppositely to *rlp44-3*, reducing phloem and xylem connectivity (Figure 3, A and B). Heterografting assays revealed that mutating *RLP44* in the rootstock reduced phloem reconnection whereas mutations in the scion non-significantly improved grafting (Supplemental Figure 3, B and C). Expression of *RLP44* was initially repressed during graft formation but within 120 hours increased higher than non-grafted controls (Figure 3C) (Melnyk et al., 2018). Genes reported to be induced by brassinosteroids such as *BR ENHANCED EXPRESSION 1* (*BEE1*), *BEE2, BEE3, PHYB ACTIVATION TAGGED SUPPRESSOR 1* (*BAS1*), *TOUCH4/XYLOGLUCAN ENDOTRANSGLUCOSYLASE/HYDROLASE 22* (*TCH4/XTH22*) *KIDARI/PRE6, SMALL AUXIN UPREGULATED-AC1* (*SAUR-AC1*), *INDOLE ACETIC ACID INDUCED 5 (IAA5*), *IAA19* and *VASCULAR RELATED NAC DOMAIN 6* (*VND6*) (Friedrichsen et al., 2002; Neff et al., 1999; Liu et al., 2020; Vert et al., 2005; Nakamura et al., 2004; Kubo et al., 2005) showed some changes in gene expression during graft healing but there was no clear or consistent pattern of up or downregulation (Supplemental Figure 3, D and E) (Melnyk et al., 2018; Zhang et al., 2022). Thus, although *RLP44* mutants showed similarities to *EVG1* mutants, these phenotypes were likely independent of canonical brassinosteroid signaling. As a second test, we analyzed callus formation levels in wounded petiole explants and observed that only *bes1-D* and *rlp44-3* significantly reduced callus formation compared to wild type Col-0 (Figure 3, E and F) and there was no clear pattern of brassinosteroid-related gene expression changes (Supplemental Figure 3F). Taken together, these results show that while core BR signaling promoted vascular regeneration and callus formation, a mutant of *RLP44* show similar regeneration phenotypes as mutations in *EVG1* and both genes appeared to repress vascular connectivity but promote callus formation.

**Figure 3.**
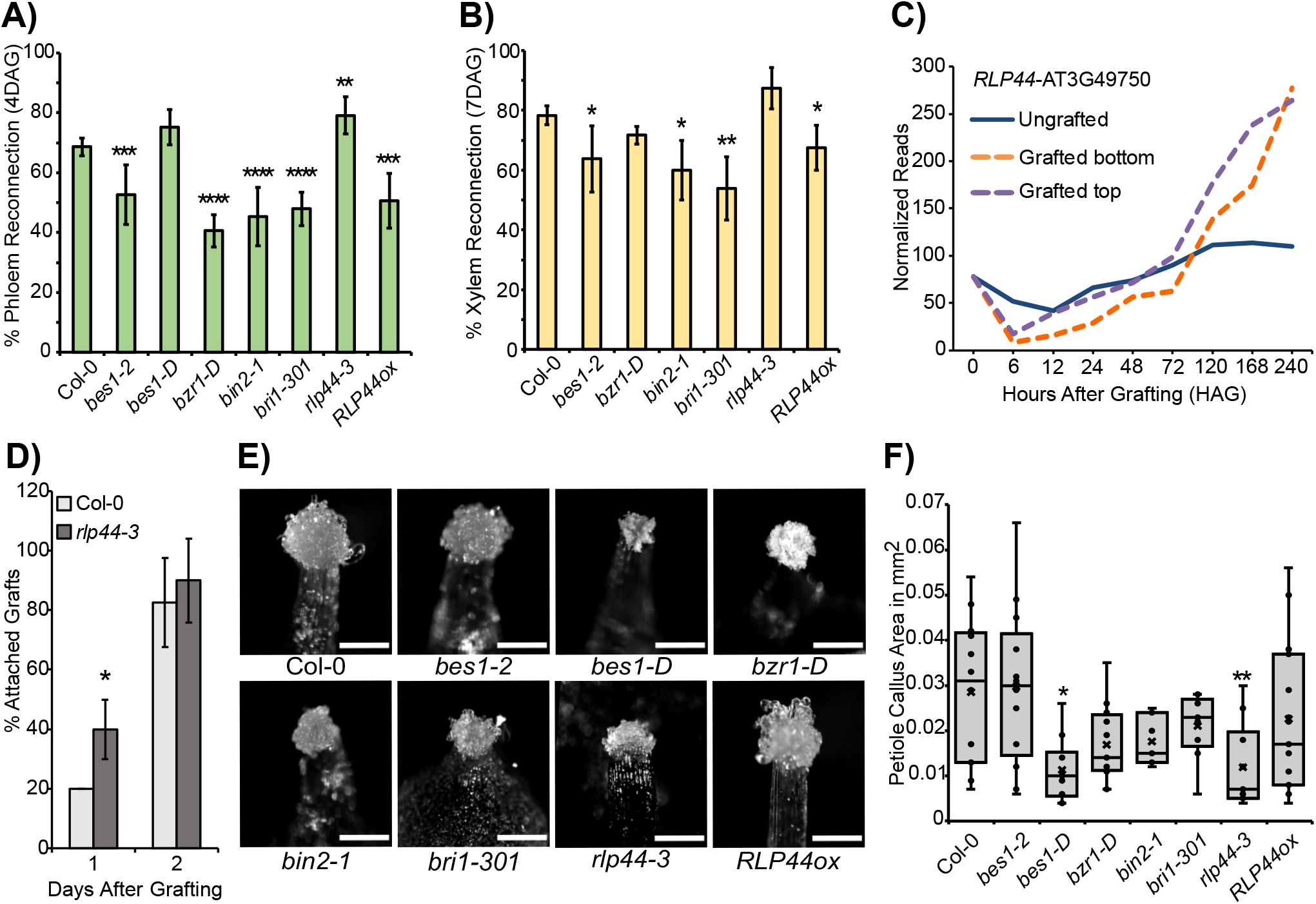
*RLP44* mutants behave like *EVG1* mutants during regeneration. (A-B) Reconnection percentage of phloem (4 days after grafting) and xylem (7 days after grafting) in homografted brassinosteroid-related mutants. Mean ± SD of 3-7 experiments with n=12 per genotype per experiment. Asterisks indicate significant difference compared to WT. * p<0.05, ** p<0.01, *** p<0.001, **** p<0.0001, pairwise t-tests with Benjamini-Hochberg adjustment. (C) *RLP44* expression during graft formation (Melnyk et al. 2018). (D) Attachment rate of Col-0 and *rlp44-3*. The mean ± SD of 3 experiments with n=12 per genotype per experiment is shown. Asterisks indicate significant differences compared to Col-0. * p<0.05, Student’s t-test. (E) Images showing callus from wounded petiole explants in brassinosteroid-related mutants, scale bars 0.2 mm. (F) Quantifications of petiole callus area in wounded petiole explants in Col-0 (n=13), *bes1-2* (n=13), *bes1-D* (n=10), *bzr1-D* (n=12), *bin2-1* (n=10), *bri1-301* (n=10), *rlp44-3* (n=10), and *RLP44ox* (n=15). Dots represent samples. Asterisks indicate significant difference compared to Col-0. * p<0.05, ** p<0.01, Wilcoxon’s test.

### *RLP44* mutants phenocopy *EVG1* mutants under non-stress conditions

To further test the links between *EVG1, RLP44* and brassinosteroids, we analyzed ectopic xylem formation using the VISUAL system and found that *rlp44-3* had enhanced ectopic xylem formation whereas *RLP44ox* showed a reduction in ectopic xylem formation when compared to wild type Col-0 (Figure 4, A and B). Brassinosteroid mutants have been previously implicated in affecting VISUAL and we found that *bes1-2* and *bin2-1* reduced ectopic xylem formation, while *bes1-D* increased ectopic xylem compared to wild type Col-0 (Supplemental Figure 4, A and B) (Kondo et al., 2015). Cross sections from *rlp44-3* roots revealed both cambium area and xylem area were reduced compared to that of wild type Col-0 (Figure 4, C-E) while *bes1-2* and *bri1-301* had no discernible reduction in cambium area compared to wild type Col-0, they did reduce xylem area. Moreover, *bin2-1* showed both reduction in cambium and xylem area when compared to wild type Col-0 (Supplemental Figure 4, C-E). The canonical brassinosteroid mutants, with the exception of *bri1-301*, had little effect on metaxylem strand number in the primary root but *rlp44-3* had extra metaxylem strands as previously reported (Figure 4F) (Holzwart et al., 2018). We then treated *rlp44-3, bri1-301, bes1-2, bes1-D, bzr1-D*, and *bin2-1* with 10nM epiBL or mock conditions. epiBL rescued the metaxylem phenotype in all mutants but *bes1-2, bri1-301*, and *rlp44-3* were resistant to xylem identity changes compared to treated wild type Col-0 (Figure 4, F and G; Supplemental Figure 4F). Overall, although some brassinosteroid mutants shared phenotypes with *evg1-1*, none except for *rlp44-3* showed complete overlap during non-stressed conditions.

**Figure 4.**
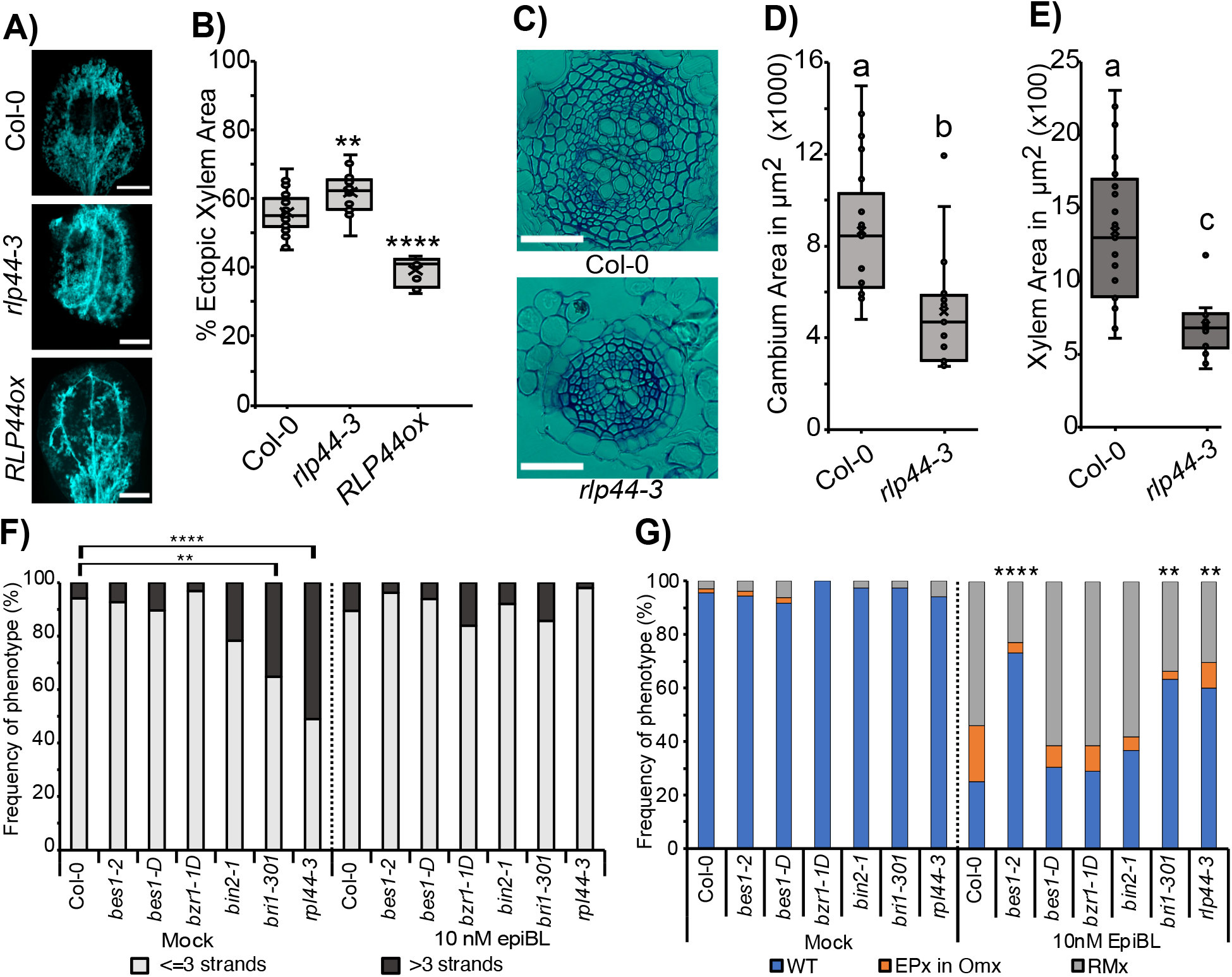
*RLP44* mutants behave like *EVG1* even in non-wounded conditions. (A) VISUAL assay images of ectopic xylem formation in *Col-0, rlp44-3* and *RLP44ox*, scale bars 1 mm. (B) Ectopic xylem quantifications of Col-0 (n=31), *rlp44-3* (n=22) and *RLP44ox* (n=10). Dots represent samples. Asterisks indicate significant difference compared to Col-0. * p<0.05, ** p<0.01, **** p<0.0001, Wilcoxon’s test. (C) Cross sections 2 mm below the shoot-root junction of Col-0 and *rlp44-3*. Scale bar represents 50μM. (D-E) Cambium area and xylem area quantifications in Col-0 (n=21) and *rlp44-3* (n=17). Dots represent individual samples. Compact letters indicate significant differences. One-way ANOVA, with Tukey’s post-hoc test. (F) Metaxylem strand number in Col-0 (n=68), *bes1-2* (n=55), *bes1-D* (n=48), *bzr1-D* (n=32), *bin 2-1* (n=37), *bri1-301* (n=37) and *rlp44-3* (n=51) under mock and epiBL treatments. Asterisks indicate significant difference compared to Col-0. ** p<0.001, **** p<0.0001, Fisher’s exact test with Benjamini-Hochberg adjustment. (G) Xylem phenotypes of Col-0 (n=68), *bes1-2* (n=55), *bes1-D* (n=48), *bzr1-D* (n=32), *bin 2-1* (n=37), *bri1-301* (n=37) and *rlp44-3* (n=51) under mock and exogenous epiBL treatments: wild type (WT), extra protoxylem in outer metaxylem (EPx in Omx) or reticulate metaxylem (RMx). Asterisks indicate significant difference compared to epiBL treated Col-0. ** p<0.001, **** p<0.0001, Fisher’s exact test with Benjamini-Hochberg adjustment.

### *EVG1* mutants affect cell wall related and ABA responsive genes

To better understand the role of *EVG1*, we performed transcriptomic analyses on *evg1-1* seedlings to identify differentially expressed genes (DEGs). We found 977 DEGs in *evg1-1* compared to wild type Col-0 with 369 genes down regulated and 608 genes up regulated (Supplemental Data Set 1). We performed a Gene Ontology analysis on *evg1-1* regulated transcripts and observed a large and significant enrichment for cellular components associated with the cell wall (Supplemental Data Set 2). Cell wall related genes such as *XYLOGLUCAN ENDOTRANSGLUCOSYLASE/HYDROLASE 19* (*XTH19*), *XTH20, XTH31*, and *TRICHOME BIREFRINGENCE-LIKE 15* (*TBL15*) were downregulated, while *XTH4, XTH6, XTH16, XTH22, XTH24, XTH27* were upregulated. Genes enriched for cell wall loosening including *EXPANSIN B3* (*EXPB3*), *EXPA3, EXPA4, EXPA5, EXPA8, EXPA15* were also downregulated (Figure 5A; Supplemental Figure 5, A and B; Supplemental Data Set 1). Our GO analysis also revealed a strong enrichment for ABA related genes in the 977 DEGs in *evg1-1* (Supplemental Data Set 2). ABA receptors such as *PYRABACTIN RESISTANCE 1* (*PYR1*) and associated paralogs such as *PYR1-LIKE 4* (*PYL4*), *PYL5* and *PYL6* were upregulated (Figure 5B). Genes related to ABA response such as *RESPONSE TO DESSICATION 29A* (*RD29A*), *KIN1, COR6*.*6, COLD RELATED 15B* (*COR15B)* were downregulated (Figure 5B, Supplemental Figure 5, C and D; Supplemental Data Set 1). Since we observed that many *evg1-1* regulated transcripts were ABA responsive, we tested whether *EVG1* itself was ABA responsive by treating *EVG1pro:GFP* with exogenously applied ABA. 3 hours post ABA treatment did not affect GFP levels, however, by 24 hours there was a reduction in GFP signals in primary roots and lateral roots compared to mock treated plants (Figure 5, C and D, Supplemental Figure 5E). Previously, it was found that exogenous application of ABA differentiates new protoxylem strands and changes the morphology of the metaxylem (Ramachandran et al., 2018). We treated *evg1-1* and *EVG1-OE* with ABA and observed that while *evg1-1* displayed no difference in xylem morphology response compared Col-0 wild type, *EVG1-OE* reduced xylem morphology changes compared to *evg1-1* (Figure 5E). When *RLP44* mutants were treated with ABA, *rlp44-3* displayed no differences in xylem morphology response compared Col-0 wild type, whereas *RLP44ox* reduced xylem morphology changes when compared to *rlp44-3* (Supplemental Figure 5F). Our results suggest that *EVG1* was negatively regulated by ABA and, like *RLP44*, the overexpressing line was ABA resistant compared to the loss of function mutant.

**Figure 5.**
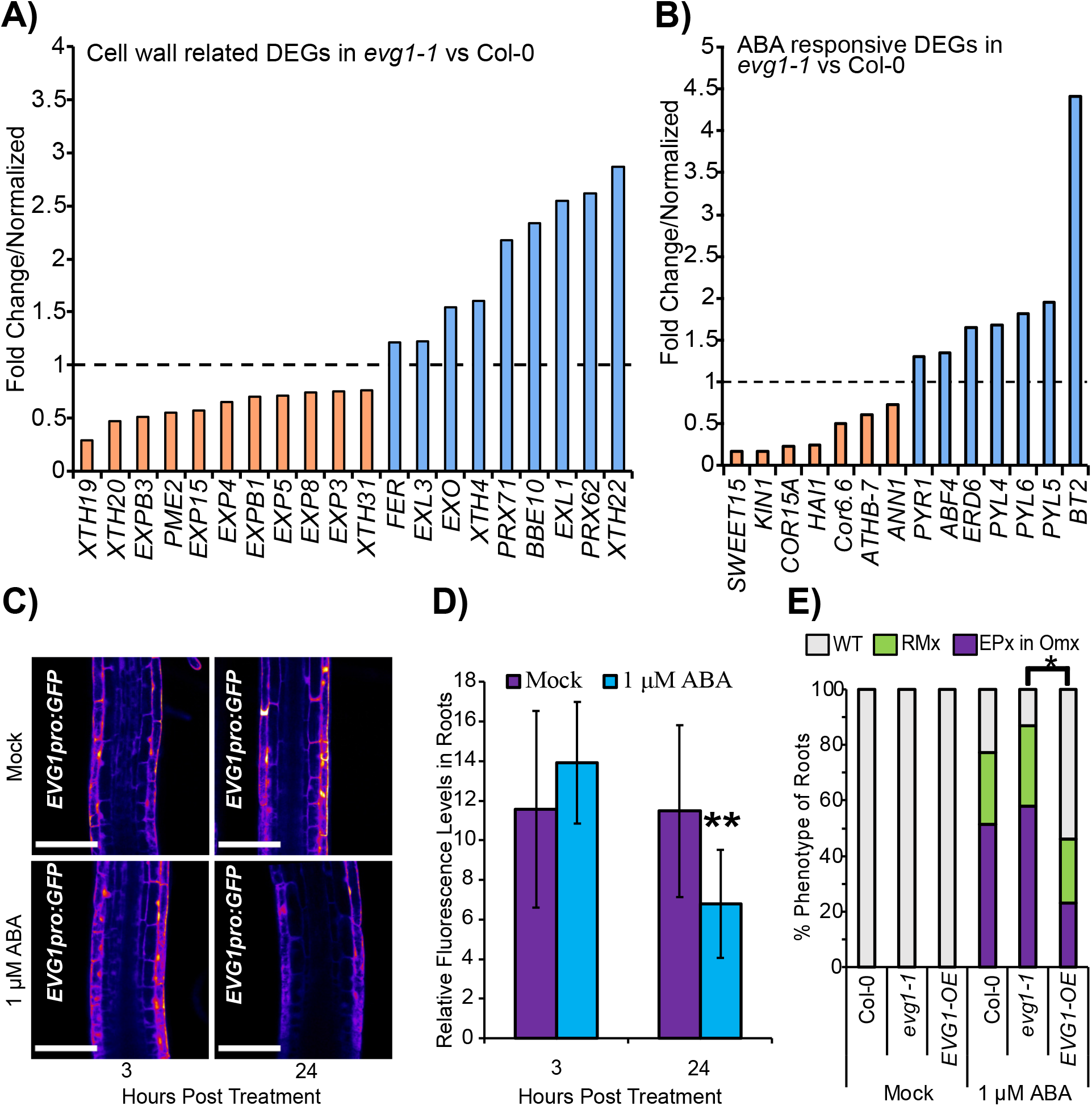
*EVG1* affects cell wall related and ABA responsive genes. (A) Cell wall related genes differentially expressed in *evg1-1*, normalised to wild type Col-0. (B) ABA response related genes differentially expressed in *evg1-1*, normalised to wild type Col-0. (C) *EVG1pro:GFP* fluorescence in the root transition zone after 3 and 24 hours of mock and ABA treatment. Scale bars represent 100 µm. (D) Relative fluorescence levels of *EVG1pro:GFP* under mock and 1 μM ABA conditions at 3 and 24 hours post treatment. Asterisks indicate significant differences between mock and treatment. ** p<0.01, Student’s t-test. (E) Xylem phenotypes of Col-0 (n=33), *evg1-1* (n=31) and *EVG1-OE* (n=13) roots with mock pr 1 µM ABA treatment. Asterisks indicate significant difference, * p<0.05, Fisher’s exact test with Benjamini-Hochberg correction.

### *EVG1* and *RLP44* regulate a common set of targets to repress regeneration

To understand how *EVG1* and *RLP44* function, we compared gene expression changes between mutants in these two genes. We performed RNA sequencing on *rlp44-3* seedlings and identified 171 DEGs, including 129 genes up regulated and 42 genes down regulated when compared to wild type Col-0 (Supplemental Data Set 3). We then compared the gene expression profiles of *evg1-1* to *rlp44-3* and observed a significant overlap between both down regulated and up regulated genes (Figure 6A). These included cell wall related genes such as *XTH19, XTH22, EXORDIUM* (*EXO*) and *EXORDIUM LIKE 3* (*EXL3*), and stress related genes such as *COLD RELATED 15B* (*COR15B*) and *RELATED TO ABI3/VP1 1* (*RAV1*) (Figure 6B). We took the 84 overlapping upregulated genes and 17 overlapping downregulated genes and looked at their expression in the grafting datasets (Melnyk et al., 2018). We observed common genes upregulated in *evg1-1* and *rlp44-3* were largely repressed in the grafted top during grafting in wild-type Col-0 (Figure 6C) whereas in grafted bottom these genes showed a more varied response (Supplemental Figure 6A). Common genes downregulated in *evg1-1* and *rlp44-3* were slightly upregulated in both grafted top and grafted bottom during grafting in wild-type Col-0 (Figure 6D; Supplemental Figure 6A). Several genes upregulated in *evg1-1* and *rlp44-3* showed upregulation in both grafted top and bottom during grafting in wild-type Col-0 including *EXORDIUM LIKE 3* (*EXL3;* AT5G51550). We wondered whether the enhanced grafting phenotypes of *evg1-1* and *rlp44-3* might be due in part to genes such as *EXL3*, so we obtained a mutant in *EXL3* and tested it in grafting and VISUAL assays. We found that loss of function *exl3* mutants reduced graft healing and reduced VISUAL ectopic xylem formation (Figure 6, E-G). These data suggest that *EVG1* and *RLP44* repressed a similar set of positive regulators of graft formation such as *EXL3*. Notably, we found *EVG1* transcripts were highly and rapidly induced upon cutting, wounding, or grafting whereas *RLP44* transcripts were not (Figure 6, H and I, Supplemental Figure 6B). In addition, *EVG1* and *RLP44* translational reporters had signal in different cell layers: EVG1 was epidermal while RLP44 was in the inner vascular tissues (Figure 6J). Thus, although these mutants shared phenotypes, we believe the genes are acting in different cellular locations and with different response dynamics to promote a common process.

**Figure 6.**
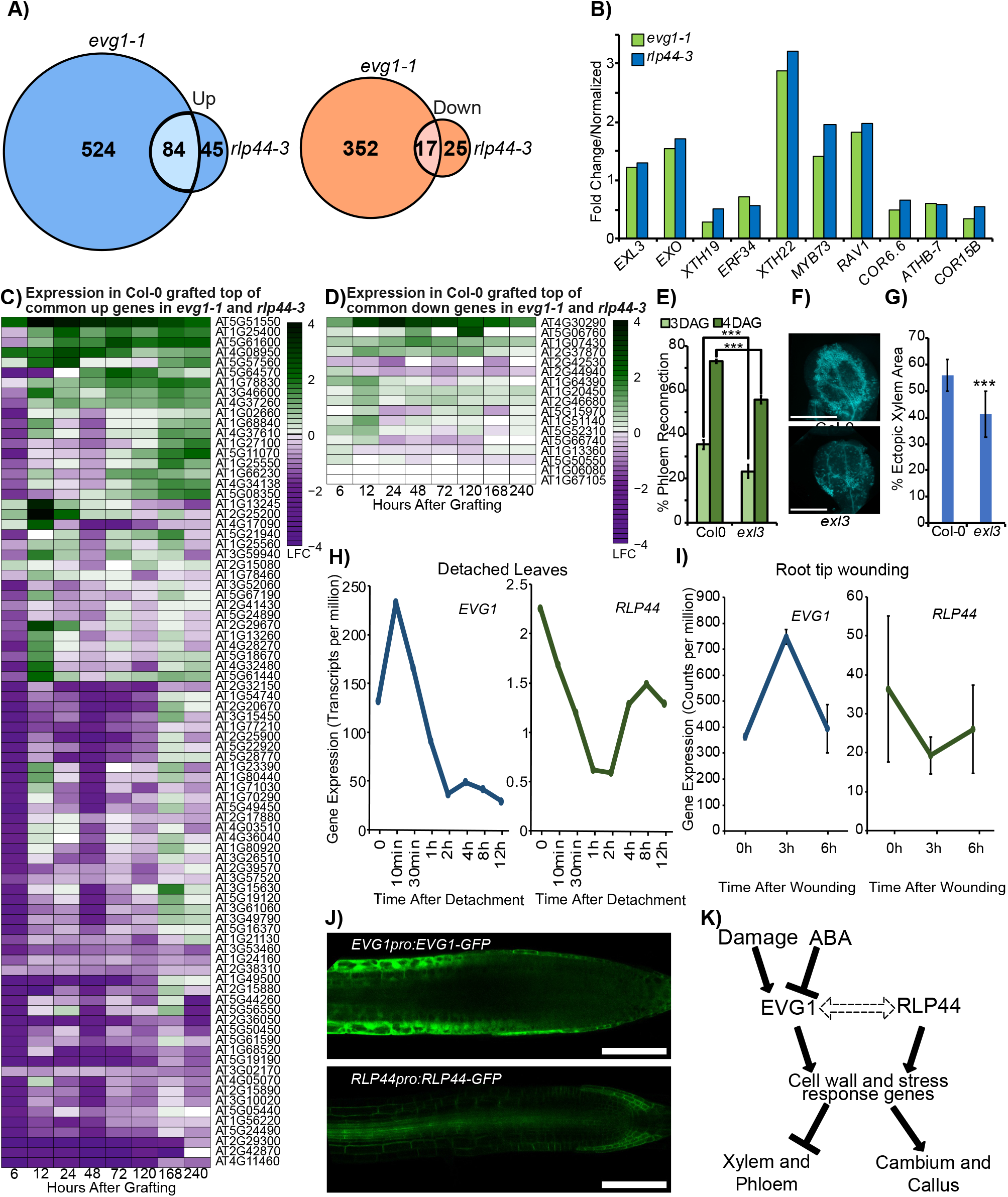
*EVG1* and *RLP44* share common targets that modify regeneration potential. (A) Venn diagram representing overlap between differentially expressed genes in *evg1-1* and *rlp44-3*. p-value 1.114E-28 for overlapping down regulated genes and 1.511E-35 for overlapping up regulated genes by hypergeometric test. (B) Fold change of a subset of genes differentially expressed in both *evg1-1* and *rlp44-3*. (C) Heat map with the 84 commonly increased transcripts from *evg1-1* and *rlp44-3* showing their expression in wild-type Col-0 grafted top tissues during graft formation compared to ungrafted tissues (Melnyk et al.,2018). (D) Heat map with the 17 commonly decreased transcripts from *evg1-1* and *rlp44-3* showing their expression in wild-type Col-0 grafted top tissues during graft formation compared to ungrafted tissues (Melnyk et al.,2018). (E) Percent phloem reconnection of Col-0 and *exl3* at 3 and 4 days after grafting (DAG). The mean ± SD of 3 experiments with n=20 per genotype per time point per experiment is shown. Asterisks indicate significant differences compared to Col-0. *** p<0.001, Student’s t-test. (F) VISUAL assay images of ectopic xylem formation in Col-0 and *exl3*. Scale bar represents 1 mm. (G) Ectopic xylem quantifications of *exl3* (n=20) and Col-0 (n=20). Asterisks indicate significant differences compared to Col-0. *** p<0.001, Student’s t-test. (H) Expression of *EVG1* and *RLP44* in 12-day old, detached leaves of seedlings (Zhang et al., 2019). (I) Expression of *EVG1* and *RLP44* in wounded root tips (Matosevich et al., 2020). (J) *EVG1pro:EVG1-GFP* and *RLP44pro:RLP44-GFP* fusion protein fluorescence. Scale bars represent 100 µm. (K) Proposed model of *EVG1* function. *EVG1* functions as a damage-activated factor that can modify downstream ABA and cell wall related genes and work together with *RLP44* to enhance callus formation but repress vascular differentiation.

## Discussion

Here, we identified *EVG1* as a stress responsive gene that repressed vascular differentiation. Upon wounding, induction of *EVG1* was rapid, occurring between 10 minutes to 6 hours after cutting (Figure 1 and 6), suggesting hormones, reactive oxygen species or control by other rapid response factors such as cell wall or turgor pressure changes (Hoermayer et al., 2020; Zhang et al., 2022; Bellandi et al., 2022). *EVG1* upregulation occurred with various biotic and abiotic stresses that caused tissue damage or tissue invasion demonstrating a common response by the plant. Unlike short-acting defence responses that happen during plant grafting (Melnyk et al., 2018), *EVG1* induction lasted for many days suggesting it had developmental roles during regeneration. *evg1* mutants reduced callus formation yet enhanced phloem connectivity, xylem connectivity and tissue attachment during wounding or grafting. To our knowledge, *evg1* is the first identified recessive mutation that enhances grafting success. These phenotypes were highly surprising given the rapid and strong induction of EVG1 by grafting, suggesting induction of a negative regulator was of benefit. *evg1* had reduced cambium levels yet enhanced grafted, suggesting an uncoupling of the apparent requirement of cambium formation on grafting success. Given the importance of cambium during grafting (Melnyk, 2017d; Melnyk et al., 2018), our results might be explained by an enhancement of differentiation in *evg1* despite limited cambium formation. To reconcile these observations, we propose that *EVG1* contributed to cambial divisions but repressed differentiation thus inhibiting processes like VISUAL, phloem reconnection and syncytium development. It likely also played a role in cell wall modifications since tissue attachment was promoted in *evg1* (Figure 1). *EVG1* also played important developmental roles under non-stress conditions since it promoted cambium formation and repressed xylem differentiation and expansion, consistent with its proposed role during regeneration. Wound-induced callus is formed from pericycle and vascular cells (Ikeuchi et al., 2017b) and genes including *EVG1* may promote cell division at the expense of cellular differentiation, thus presenting an early step in plant damage healing to seal wounds with undifferentiated cell masses.

The molecular function of EVG1 remains elusive but *in silico* analyses have found two Domain of Unknown Function 642 (DUF642) in the EVG1 protein structure (Vázquez-Lobo et al., 2012b). DUF642 is a highly conserved protein family in spermatophytes involved in cell wall modification, cell wall maintenance (Salazar-Iribe et al., 2016; Cruz-Valderrama et al., 2019) and, in *Amaranthus*, abiotic stress response (Palmeros-Suárez et al., 2017). *EVG1* has also been found in cell wall proteomes and interacts with cellulose and hemicellulose *in vitro* (Ndimba et al., 2003; Borner et al., 2003; Vázquez-Lobo et al., 2012b). Our own analyses suggest that the EVG1 protein is localized in the cell membrane or extracellular space. *in silico* analyses have also revealed possible relevant EVG1 features: it is highly coiled, compact, contains a transmembrane domain, contains a signal peptide motif at the N terminus and is predicted to be located in the extracellular space (Supplemental Figure 6, C-E; Supplemental Data Set 4) (Krogh et al., 2001; Jumper et al., 2021; Teufel et al., 2022; Varadi et al., 2022). Such a transmembrane domain may be relevant for insertion in the plasma membrane and suggests a possible role in signal transduction or signal perception. Analyses of differentially expressed genes in *evg1* revealed many related to cell wall organization, biogenesis, and cell wall loosening. X*TH4*, which affects xylem cell expansion and secondary cell wall development (Kushwah et al., 2020), was up regulated when *EVG1* function was lost. Multiple expansins such as *EXPA4, EXPA8, EXPA16, EXPB3* were also down regulated in *evg1*. EVG1 thus appears to be closely linked to cell wall function and cell wall signaling. Previous analyses found *EVG1* mRNA highly expressed in developing seeds and it’s possible that *EVG1* is relevant for aspects of germination such as pectin-related mucilage formation (Vázquez-Lobo et al., 2012b; Garza-Caligaris et al., 2012). Further work is needed to understand EVG1’s precise molecular function and its function in the cell wall including interactions with pectin and cellulose.

Although our analysis did not identify *EVG1* activating substances, we found that ABA downregulated *EVG1* and many ABA-related genes were differentially expressed in *evg1* mutants. As a stress hormone, ABA inhibits various aspects of growth including cell wall loosening, cell elongation, cell cycle and seed dormancy (Gimeno-Gilles et al., 2009; Takatsuka and Umeda, 2019; Liu et al., 2021). Low levels of ABA also induce the formation of xylem cells in *Arabidopsis* roots (Ramachandran et al., 2018). Thus, ABA might repress cell divisions and promote xylem differentiation in part via the inhibition of *EVG1*. Our analyses also uncovered a novel role for canonical and non-canonical brassinosteroid signaling during regeneration. Although we saw little evidence for a consistent brassinosteroid response at the graft junction, many brassinosteroid genes were needed for efficient grafting success (Figure 3). *RLP44* was exceptional though since it repressed grafting success. Previously this gene has been implicated in promoting cambium and repressing xylem formation (Wolf et al., 2014; Holzwart et al., 2018) and here we extend these observations and demonstrate that *rlp44* mutations enhanced vascular regeneration and graft formation. Although *RLP44* and *EVG1* mutants were phenotypically indistinguishable in our assays, notable differences were present. We saw little evidence that *RLP44* transcription was induced early during stress and the cellular localization of *RLP44* was primarily in the vascular cylinder whereas *EVG1* was primarily in the epidermis and cortex (Figure 6). Thus, we propose that these two genes share a similar signaling pathway but their localization differences suggest a mobile signal or factor links the proteins to control cambial proliferation and vascular differentiation. The transcriptional overlap between *evg1* and *rlp44* identified numerous genes regulated in common including *EXL3* and *EXO. EXO* and *EXL1*, close homologs of *EXL3*, affects cell wall expansion and development in leaves (Schröder et al., 2009). Such overlapping genes may present factors that distinguish cell division from differentiation and deserve further investigation. In summary, we propose a mechanistic model in which *EVG1* responds to stress and, together with *RLP44*, mediates cell wall signaling to activate cambium proliferation (Figure 6K). Such a framework could help modify grafting success and the regenerative abilities of plants.

## Material and Methods

### Plant material and growth conditions

*Arabidopsis thaliana* accession Columbia-0 (Col-0) was used as the wild type control in this study and all mutants used were in Col-0 background, unless mentioned otherwise. List of mutants used in this study are described in Supplemental Table 1. Primers used for checking homozygosity or for transcript quantification are described in Supplemental Table 2 and 3. Seeds were surface sterilized with 75% (v/v) ethanol for 20 minutes, then 99.5% (v/v) ethanol for 5 minutes. The seeds were put in a sterile hood to remove residual ethanol. Sterilized seeds were then placed on half-strength Murashige and Skoog (MS) (Murashige and Skoog, 1962) medium with 1.2% plant agar unless mentioned otherwise. Seeds stratified for 48 – 72 hours in 4°C were moved to the growth chamber under short day conditions (8 hours light/16 hours dark, ∼110 mmol m^-2^s^-1^, 20°C, Conviron A1000 chamber) or long day conditions 16 hours light/8 hours dark, 120 mmol m^-2^s^-1^, 22°C day temperature and 20°C night temperature) unless mentioned otherwise. Plates were kept vertically for vertical plant growth.

### Plasmid construction and transgenic line generation

To generate *EVG1pro:GFP* and *EVG1pro:EVG1-GFP*, a 2112 bp promoter region as described in (Salazar-Iribe et al., 2016), and *EVG1* coding sequence without stop codon was cloned in to the promoter module (A-B overhang) and CDS module (C-D overhang) respectively in the GreenGate cloning system (Lampropoulos et al., 2013). Following the cloning protocol all necessary modules including the GFP coding sequence module (C-D overhang) for *EVG1pro:GFP* and GFP linker sequence (D-E overhang) module were used in the final cloning reaction to create *EVG1pro:GFP* and *EVG1pro:EVG1-GFP* respectively. The module for selection was obtained from pHDE-35S-Cas9-mCherry-UBQ which was a gift from Yunde Zhao (Addgene plasmid # 78932; http://n2t.net/addgene:78932; RRID:Addgene_78932) (Gao et al., 2016). Transgenic lines were generated using the floral dip method (Clough and Bent, 1998). All primers used for cloning are listed in Supplemental Table 4.

### Ectopic xylem formation in cotyledons

Ectopic xylem formation assays were performed according to a previously published method Vascular cell Induction culture System Using Arabidopsis Leaves (VISUAL) (Kondo et al., 2015, 2016) with one minor change in the induction media with the addition of BF-170 (Nurani et al., 2020) a lignin binding secondary cell wall indicator for easier imaging of xylem. In brief, *Arabidopsis* seeds were grown for 6 days under 24 hours light conditions. The cotyledons were then excised and transferred to induction media. At the end of the 4-day induction period, cotyledons were fixed overnight in a solution of acetic acid and 99.5% ethanol (1:3, v:v). Samples were then placed in a chloral hydrate solution and mounted on slides with chloral hydrate for visualization of autofluorescence (UV filter) with a Leica M205 FA stereo fluorescent microscope. The area of ectopic xylem was calculated from autofluorescence levels using Fiji and normalized to the total cotyledon area. Cotyledon veins were excluded from the quantification.

### Plant micrografting and attachment

7-day-old short day grown seedlings were used for micro-grafting (Melnyk, 2017b). In brief, for attachment assays, grafted plants were picked up with forceps at the root-hypocotyl junction and placed back down at 1 and 2 days after grafting. If the scion remained attached during the entire movement, the plant was scored as positive for attachment. Percent attached grafts was calculated as a function of number of attached grafts to the total grafted plants. For the phloem reconnection assay, the cotyledon was damaged with forceps and carboxyfluorescein diacetate (CFDA) was placed on the wound site. Phloem reconnection was scored successful if the fluorescent signal appeared in the root after 1 hour at tested time points. Percent phloem reconnection was calculated as a function of number of plants with fluorescent roots vs number of plants grafted. New plants were used for each time point. For the xylem reconnection assay, the root 1-2cm below the hypocotyl cut and then CDFA was dropped on the wound site. After 20 minutes, xylem reconnection was score successful if the fluorescent signal was found in the cotyledon at tested time points. Percent xylem reconnection was calculated as a function of number of plants with fluorescent cotyledons vs number of plants grafted. New plants were used for each time point.

### Callus regeneration and wounding assays

Callus induction in petiole explants was performed using a previously published method with some changes (Iwase et al., 2017). Cotyledons with petioles were excised from 10-day old, long day grown seedlings. They were then placed on full strength MS medium plates supplemented with 1% sucrose and 0.6% Gelrite under long day conditions. Callus induction in hypocotyl explants was performed using a previously published method with some changes (Iwase et al., 2011b). The seeds were grown in the dark to generate etiolation on MS medium supplemented with 0.05% MES, 0.5% Sucrose, 0.8% Gelrite. 7 days of dark growth was followed with a cut that was performed at approximately 7mm above the hypocotyl-root junction to induce callus. After 8 days induction, sample tissues from both petiole explants and hypocotyl explants were imaged with a Leica M205 FA stereo fluorescent microscope. Projected callus area in the image was measured using the freehand tool in Fiji.

### Histological sections

To avoid lateral roots, 21-day old long-day grown seedling samples were cut 2 cm below the shoot tip and were collected and vacuum-infiltrated using a fixation solution (1% glutaraldehyde, 4% formaldehyde, 0.05M sodium phosphate). After keeping in the fixation solution for at least overnight and subsequent ethanol dehydration, the samples were oriented with shoot pointing to the top in a mold, with the leaves removed. The samples were then infiltrated and embedded with Leica Historesin. 2.5-µm thin cross sections 2 mm below the shoot-root junction were cut with a Leica microtome and followed by staining with Toluidine blue and imaged with Zeiss Axioscope A1 microscope.

### Confocal Microscopy and treatments

For confocal microscopy, roots were mounted in 10 μM propidium iodide (PI) solution between two coverslips and imaged immediately. Confocal micrographs were captured using Zeiss LSM780 inverted Axio Observer with supersensitive GaAsP detectors for *EVG1pro:GFP* and Zeiss LSM800 for *EVG1pro:EVG1-GFP* and *RLP44pro:RLP44-GFP* respectively. For reporter lines expressing GFP and stained with PI, 488nm excitation and 500-553nm emission was used for both GFP and PI signals. For analysis of fluorescence during grafting, grafted plants were mounted on water between two cover slips analyzed 24 hours after grafting. For wounding assays, wounded plants were mounted on water between two cover slips analyzed 24 hours after wounding. To analyze fluorescence changes after ABA treatment, images of ABA treated plants were compared to images of mock treated roots, and mean gray values were compared calculated using Fiji.

### Root xylem architecture

Sterilized and stratified seeds were placed on 25 mm pore Sefar Nitex 03–25/19 mesh (Ramachandran et al., 2018) on a half-strength MS plate supplemented with 1.2% plant agar, and then grown vertically in long day conditions for 3 days. Post 3 days the plants were transferred to half-strength MS, 1.2% plant agar plates supplemented with either 95% ethanol, DMSO, 1 μM ABA or 10nM epiBrassinolide and kept vertically in long day conditions for additional 3 days. Roots were mounted on chloral hydrate solution and imaged under 40x with Zeiss Axioscope A1 with differential interference contrast (DIC) to analyze xylem morphology.

### RNA isolation and quantitative real time polymerase chain reaction

Total RNA was isolated using a Roti-Prep RNA MINI Kit. RNA samples were quantified using a NanoDrop ND-1000 spectrometer (Thermo Fisher Scientific). cDNA was prepared using 500 ng of total RNA using Maxima First Strand cDNA Synthesis Kit containing oligo(dT) and random hexamer primers. The cDNA was diluted 1:9 with nuclease-free water. iCycler iQ Real-Time PCR detection system with 10 μL reaction volumes (5 μL of 2X Maxima SYBR Green qPCR/ROX Master Mix, 1.2 μM of forward and reverse primers, and 2.5 μL of diluted cDNA) was used to perform the qPCR. The program used for qRT-PCR was as follows: initial denaturation for 10 minutes at 95°C followed by 40 cycles of 95 °C for 30 seconds, 60 °C for 30 seconds. This was followed by a melt curve analysis. Relative expression levels of selected genes were calculated using 2^-ΔΔCT^ method (Livak and Schmittgen, 2001). For analysis of *evg1-1* and *evg1-2* lines *UBC9, TIP41-like, PP2A* were used as loading reference (Czechowski et al., 2005). For analysis of *EVG1* transcript levels during VISUAL, *APT1* was used as a loading reference (Gutierrez et al., 2008). Three biological replicates were prepared for each genotype.

### Preparation, sequencing and analysis of transcriptomic library

For RNAseq library preparation, 200ng of total RNA extracted from seven-day old, short-day grown seedlings was treated using a Poly(A) mRNA Magnetic Isolation Module kit. The library was prepared with the resulting mRNA using a NEBNext® Ultra(tm) II Directional RNA Library Prep Kit for Illumina® and NEBNext Multiplex Oligos for Illumina. Libraries were sequenced at Novogene on a NovaSeq 6000 in 150bp paired-end mode. For RNAseq analyses, the raw data were cleaned using fastp to remove the low-quality reads (Chen et al., 2018). Hisat2 was used to map the cleaned reads to the Arabidopsis reference TAIR10 (Kim et al., 2015). Counts of reads were determined using HTseq-count (Anders et al., 2015). Differentially expressed genes (DEG) were defined using the DESeq2 R package. Genes with an adjusted p-value < 0.05 were considered to have statistically significant expression differences between samples, wild type Col-0 was the reference. The list of DEGs between *evg1-1* vs Col-0 is provided in Supplemental Data Set 1. GO term enrichment analysis for *evg1-1* DEGs was performed using the GO term enrichment tool on TAIR relying on PANTHER (Mi and Thomas, 2009; Mi et al., 2013, 2019, 2021; Ashburner et al., 2000; The Gene Ontology Consortium et al., 2021) and is provided in Supplemental Data Set 2. The list of DEGs between *rlp44-3* vs Col-0 is provided in Supplemental Data Set 3.

### Nematode infection assays

The nematode infection was performed following the protocol mentioned in previous reports (Anjam et al., 2020). Briefly, 12 days old Arabidopsis plants grown on modified Knop medium were infected with approximately 100 freshly hatched surface-sterilized second stage juveniles (J2s) *Heterodera schachtii*. After 12 days post infection (dpi), developed male and female nematodes were counted using a Leica MZ16 stereo zoom microscope. At 14 dpi, syncytia and females were imaged using a Leica MZ16 stereo zoom microscope mounted with Leica MC190HD Camera. The area of the corresponding images was measured using Image J.

### Statistical analyses

All statistical analyses were performed using R Studio with R version 4.2.0. Student’s t-test with two tail distribution was used to compare two groups in case of normal distribution, otherwise Wilcoxon’s signed rank test was used. For categorical values, Fisher’s exact test with Benjamini-Hochberg correction was used. For comparison between multiple groups, One-way ANOVA followed by a post-hoc Tukey HSD test was performed. The p values <0.05 were considered statistically significant.

## Accession numbers

Sequence data from this article can be found in the EMBL/GenBank data libraries under the following accession numbers: *EVG1* (AT3G08030), *CIPK5* (AT5G10930), *LTPG5* (AT3G22600), *SVB5* (AT4G24130), *RLP44* (AT3G49750), *EXL3* (AT5G51550), *BRI1* (AT4G39400), *BES1* (AT1G19350), *BZR1* (AT1G75080), *BIN2* (AT4G18710). mRNA sequencing data from this study were deposited in the Gene Expression Omnibus database https://www.ncbi.nlm.nih.gov/geo under accession number GSE224565.

## Supplemental Data

Supplemental Figure S1: Expression profiles of *EVG1* during stress, nematode infection, grafting, and wounding.

Supplemental Figure S2: *EVG1* expression levels and phenotypes.

Supplemental Figure S3: Brassinosteroid responses and requirements at the graft junction.

Supplemental Figure S4: Brassinosteroid mutants affect VISUAL and vascular development.

Supplemental Figure S5: ABA and cell wall effects in *evg1*

Supplemental Figure S6: EVG1 is stress responsive and possesses a signal peptide and a transmembrane domain.

Supplemental Data Set 1: List of DEGs in *evg1-1* compared to Col-0.

Supplemental Data Set 2: GO term enrichment analysis for DEGs in *evg1-1*.

Supplemental Data Set 3: List of DEGs in *rlp44-3* compared to Col-0.

Supplemental Data Set 4: Probability and prediction scores of amino acids for signal peptide and transmembrane domain.

Supplemental Table 1: List of genotypes used in this study

Supplemental Table 2: Primers used for genotyping

Supplemental Table 3: Primers used for expression analysis

Supplemental Table 4: Primers used for cloning

## Acknowledgements

We thank the Nottingham Arabidopsis Seed Center (NASC), Sebastian Wolf (University of Tübingen, Germany), Yuki Kondo (Kobe University, Japan), and RIKEN BRC for materials. We would also like to thank Igor Sabljic (Swedish University of Agricultural Sciences) for help with protein prediction software and Abdul Kareem V.K (Swedish University of Agricultural Sciences) for confocal microscopy assistance. S.M. and C.W.M were supported by a Vetenskapsrådet grant (2017-05122). A.Z. and C.W.M were supported by a Wallenberg Academy Fellowship (2016-0274). C.M. and C.W.M were supported by a European Research Council starting grant (GRASP-805094). M.S.A. and P.M. were supported by a Vetenskapsrådet grant (2019-05634) and a MSCA Postdoctoral Fellowship (101066035-PREENER).

## Competing interests

None declared.

## Author contributions

S.M. and C.W.M designed the study. S.M. conducted the experiments, analyzed the data, and designed the figures. A.Z. performed the transcriptomic analyses. C.M. performed the histological sections, C.M. and S.M. analyzed the data. M.S.A. and P.M. performed and analyzed nematode infection data. S.M. and C.W.M. wrote the manuscript. All authors approved the final manuscript. Funding acquisition by A.Z., P.M. and C.W.M.

## Data availability

All data is available on request from the corresponding author.

## Notes

### Competing Interest Statement

The authors have declared no competing interest.

