## Supplemental Figures 1-6 for "Damage activates *EVG1* to suppress vascular differentiation during regeneration in *Arabidopsis thaliana*"

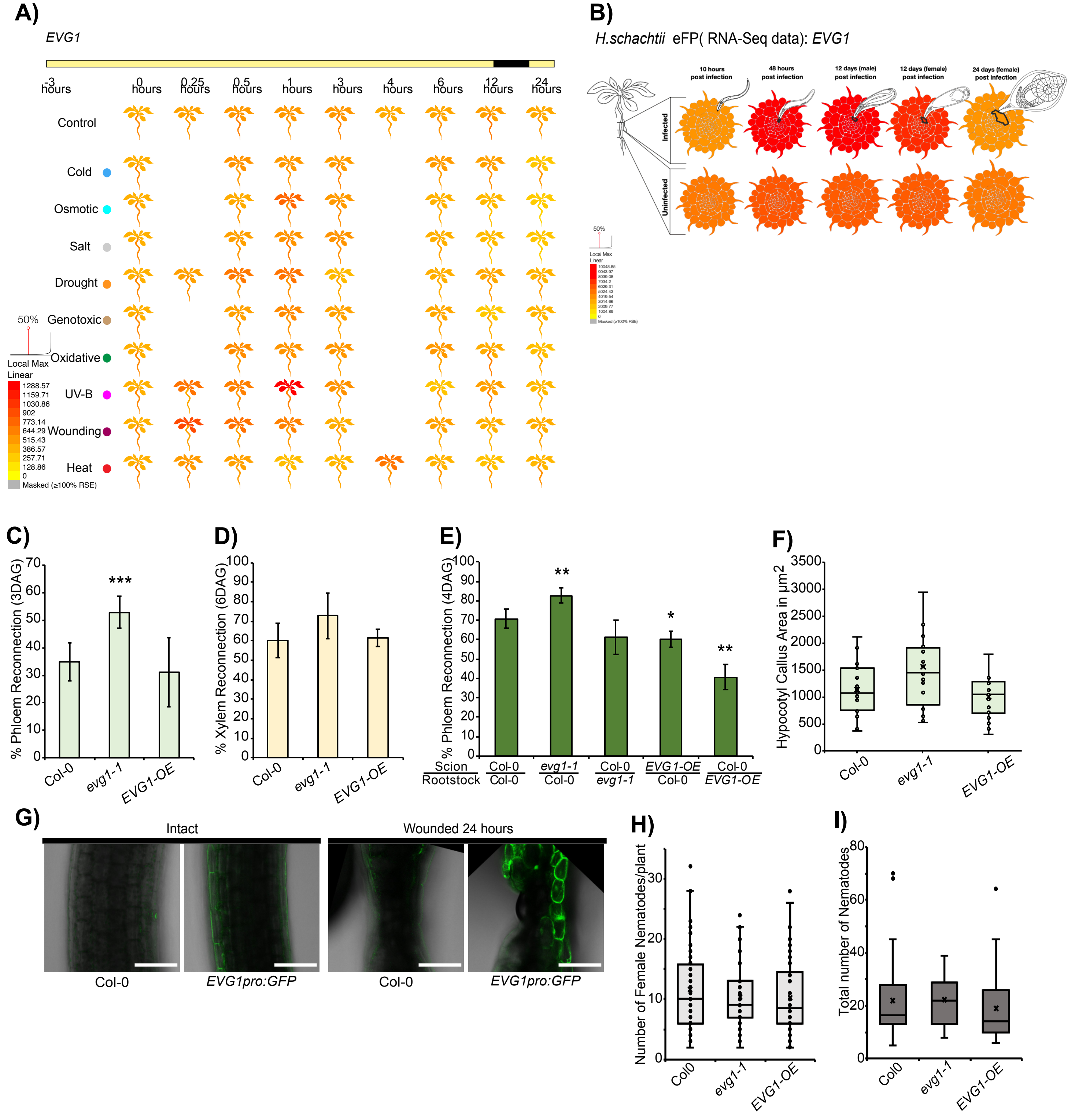

**Supplemental Figure 1. *Expression profiles of EVG1* during stress, nematode infection, grafting and wounding.** (A) *EVG1* expression during different abiotic stresses. Data from ePlant browser (Bar Toronto) (Kilian et al., 2007; Fucile et al., 2011; Waese et al., 2017). (B) *EVG1* expression during *H. schachtii* infection. Data from ePlant browser (Bar Toronto) (Fucile et al., 2011; Waese et al., 2017). (C) Percent phloem reconnection of Col-0, *evg1-1* and *EVG1-OE* 3 days after grafting (DAG). The mean ± SD of 5 experiments with n=12 per genotype per experiment is shown. Asterisks indicate significant differences compared to Col-0. *** p<0.001, Student's t-test. (D) Percent xylem reconnection of Col-0, *evg1-1* and *EVG1-OE* 6 DAG compared to Col-0. The mean ± SD of 4 experiments with n=12 per genotype per experiment is shown. (E) Percent phloem reconnection of *EVG1* heterografting at 4DAG. The mean ± SD of 4 experiments with n=12 per genotype per experiment is shown. Asterisks indicate significant differences compared to Col-0. * p<0.05, ** p<0.01, Student's t-test. (F) Hypocotyl explant callus areas of Col-0, *evg1-1* and *EVG1-OE*. Dots represent individual samples (n=25 per genotype). (G) *EVG1pro:GFP* fluorescence at the hypocotyl wound site 24 hours after wounding, scale bars, 100 μm. (H) Average number of female nematodes per plant in Col-0, *evg1-1* and *EVG1-OE* 12 days post infection. (I) Box plot indicating total number of nematodes per plant in Col-0, *evg1-1* and *EVG1-OE*. Dots indicate individual samples.

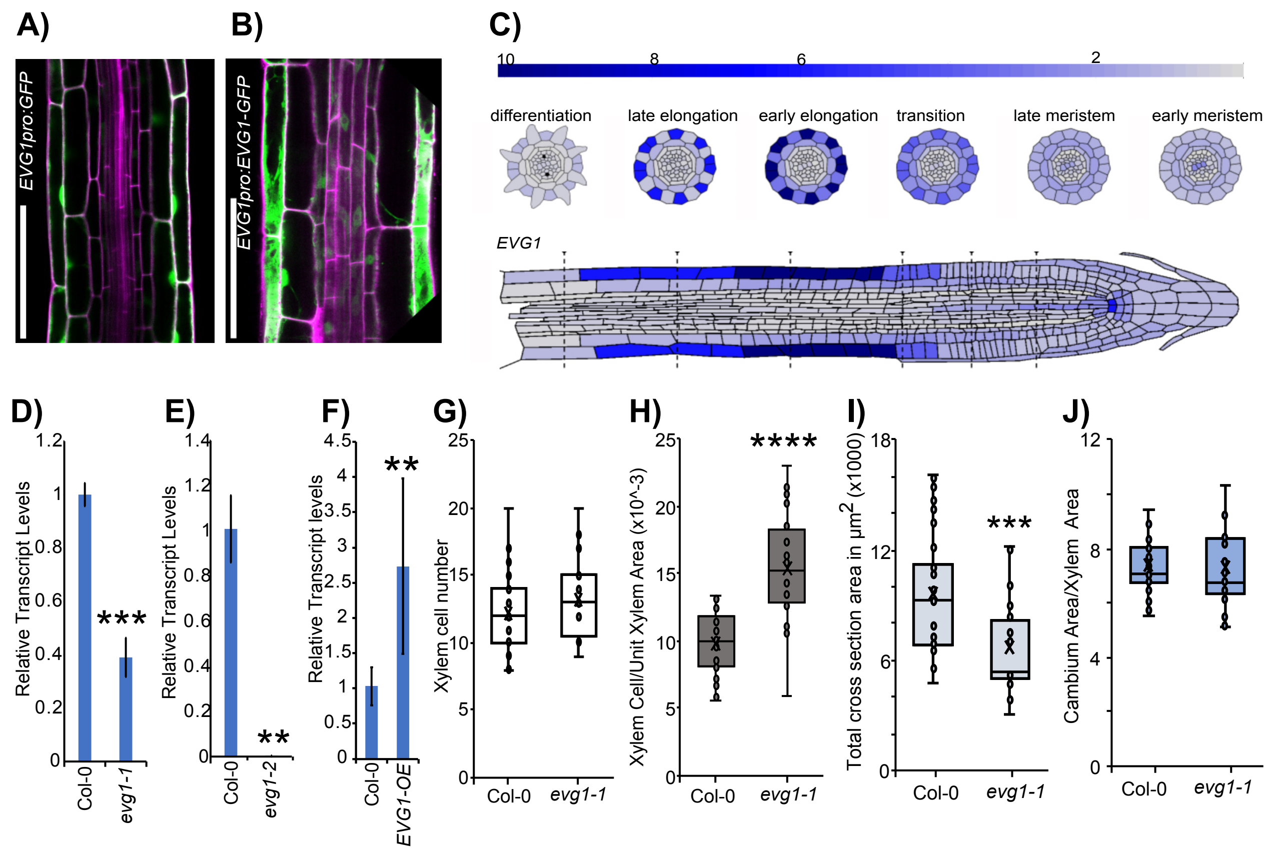


**Supplemental Figure 2. *EVG1* expression levels and phenotypes.** (A) *EVG1pro:GFP* fluorescence at the transition zone in the roots. Cell walls stained by PI (magenta). Scale bar represents 100 µm. (B) *EVG1pro:EVG1-GFP* fluorescence at the transition zone in the roots. Cell walls stained by PI (magenta). Scale bar represents 100 µm. (C) *EVG1* expression pattern in different cell layers of the primary root as generated from Root Cell Atlas (https://rootcellatlas.org/)Ryu et al., 2019; Denyer et al., 2019; Shulse et al., 2019; Jean-Baptiste et al., 2019; Wendrich et al., 2020; Shahan et al., 2022). (D-E) Expression of *EVG1* in *evg1-1* and *evg1-2* T-DNA insertion lines. Asterisks indicate significant difference in transcript levels of *EVG1* between Col-0 and mutant. ** p <0.01, *** p <0.001, Student's t-test. (F) Expression of *EVG1* in FOX hunting line *EVG1-OE.* Asterisks indicate significant difference in transcript levels of *EVG1* between Col-0 and mutant. **, p <0.01, Student's t-test. (G-J) Number of xylem cells, xylem cell/unit xylem area, total cross section area and cambium:xylem area of 21-day old Col-0 and *evg1-1* shoot-root cross sections. Dots represent individuals Col-0 (n=35), *evg1-1* (n=24). Asterisk show statistical significance compared to Col-0. *** p<0.001****, p<0.0001 Wilcoxon's test.

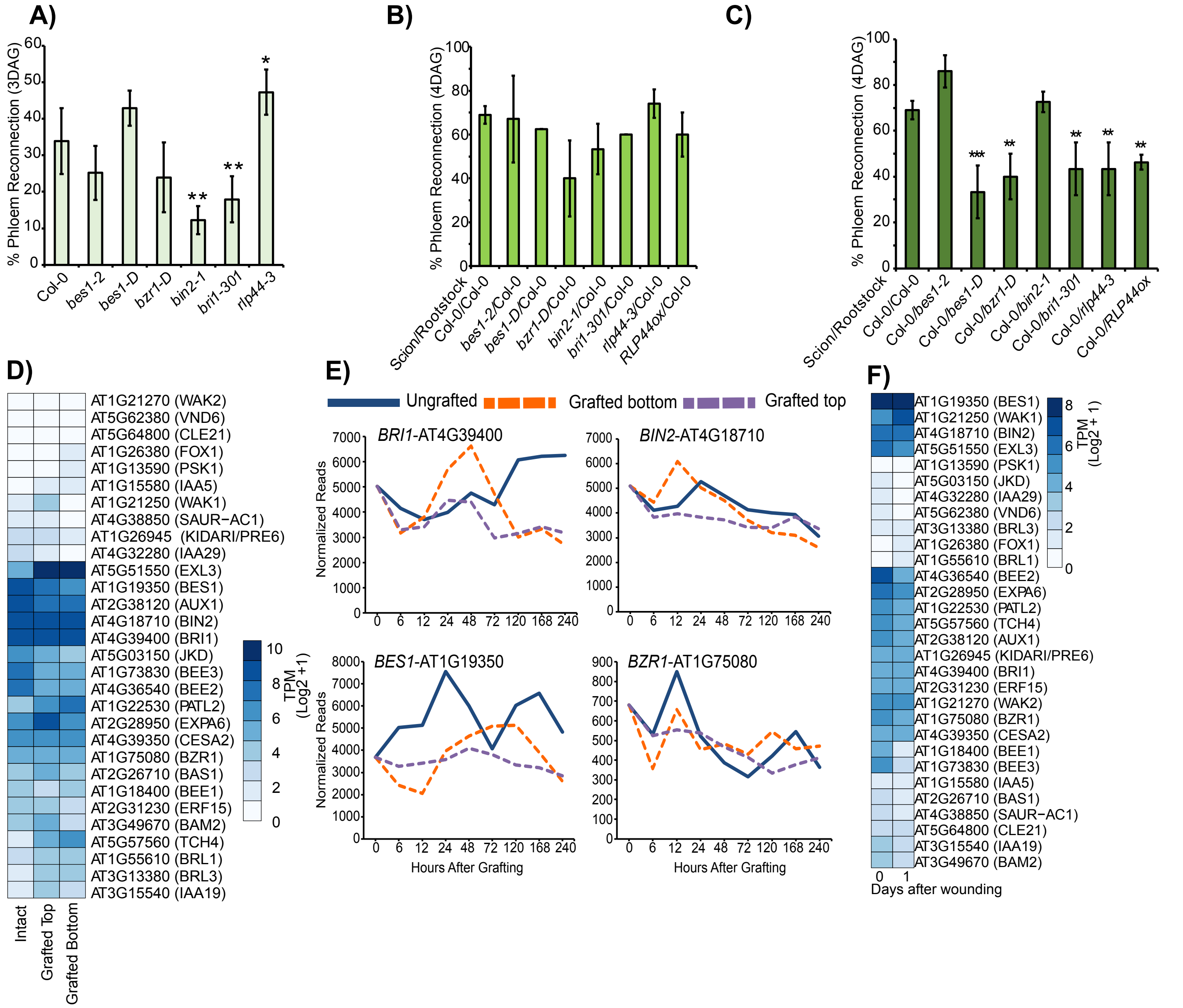


**Supplemental Figure 3: Brassinosteroid responses and requirements at the graft junction.** (A) Percent phloem reconnection of brassinosteroid signaling mutants three days after grafting (DAG) compared to WT. Mean ± SD of 4 experiments with n=12 per genotype per experiment. Asterisks indicate significant differences compared to wild-type Col-0. * $p<0.05$, ** $p<0.01$, pairwise t-tests with Benjamini-Hochberg adjustment. (B-C) Percent phloem reconnection during heterografting at 4 DAG of brassinosteroid signaling mutants as rootstocks or scions. Mean ± SD of 3 experiments with n=12 per genotype per experiment. Asterisks indicate significant differences compared to wild-type Col-0. ** $p<0.01$, *** $p<0.001$, pairwise t-tests with Benjamini-Hochberg adjustment. (D) Heat map of the expression levels of brassinosteroid responsive genes at 24 hours after grafting (Zhang et al., 2022). (E) Expression profile of brassinosteroid induced genes during graft formation (Melnyk et al. 2018). (F) Heat map of the expression levels of brassinosteroid responsive genes in petioles after wounding (Pan et al., 2019).

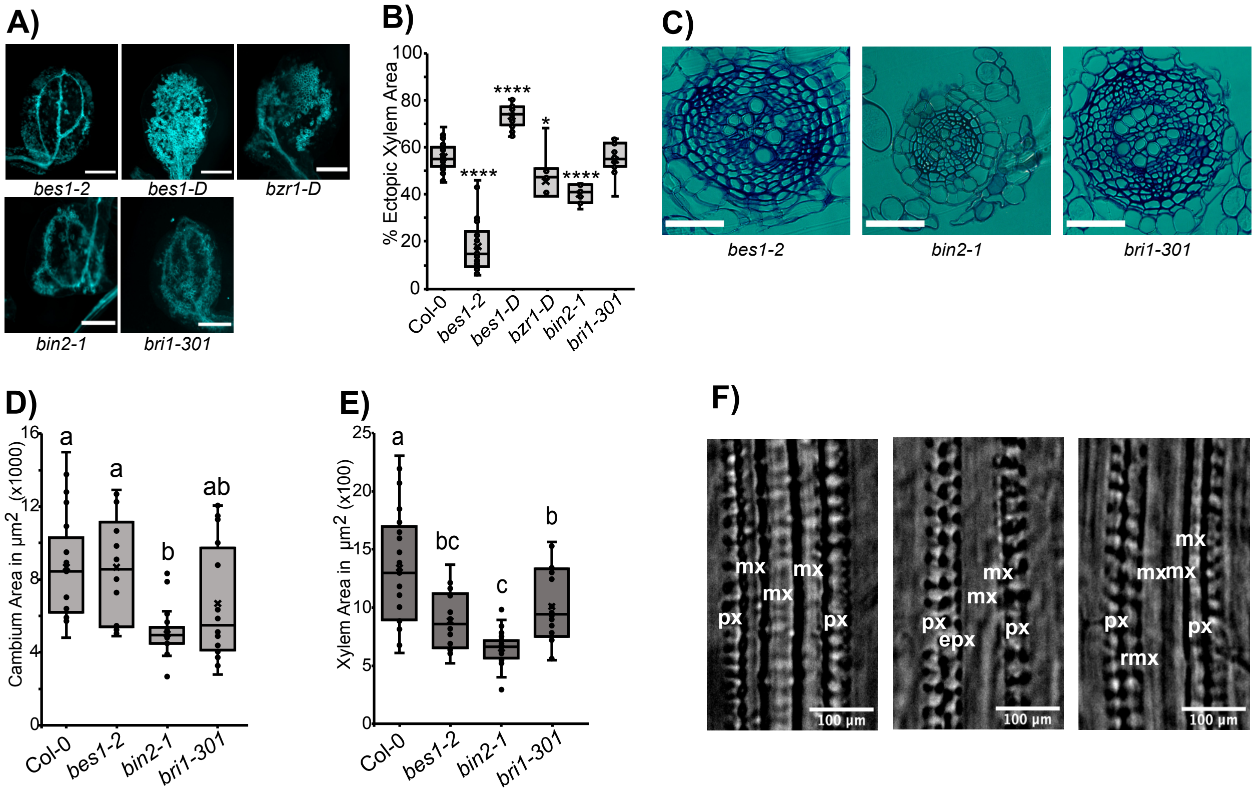


**Supplemental Figure 4. Brassinosteroid mutants affect VISUAL and vascular development.** (A) VISUAL assay images of ectopic xylem formation in *Col-0, bes1-2. bes1-D, bzr1-D, bin2-1* and *bri1-301*. Scale bars 1 mm. (B) Ectopic xylem quantifications of Col-0 (n=31), *bes1-2* (n=22), *bes1-D* (n=22), *bzr1-D* (n=10), *bin2-1* (n=10) and *bri1-301* (n=15). Dots represent samples. Asterisks indicate significant difference compared to Col-0. * p<0.05, ** p<0.01, **** p<0.0001, Wilcoxon's test. (C) Cross sections 2 mm below the shoot-root junction of *bes1-2*, *bin2-1*, *bri1-301* and *rlp44-3*. Scale bar represents 50µM. (D-E) Cambium area and xylem area in Col-0 (n=21), *bes1-2* (n=14), *bin2-1* (n=27), and *bri1-301* (n=20). Dots represent individual samples. Compact letters indicate significant differences. One-way ANOVA, with Tukey's post-hoc test. (F) Images showing primary root xylem phenotypes including protoxylem (px), metaxylem (mx), extra protoxylem (epx), reticulate metaxylem (rmx). Scale bar represents 100µM.

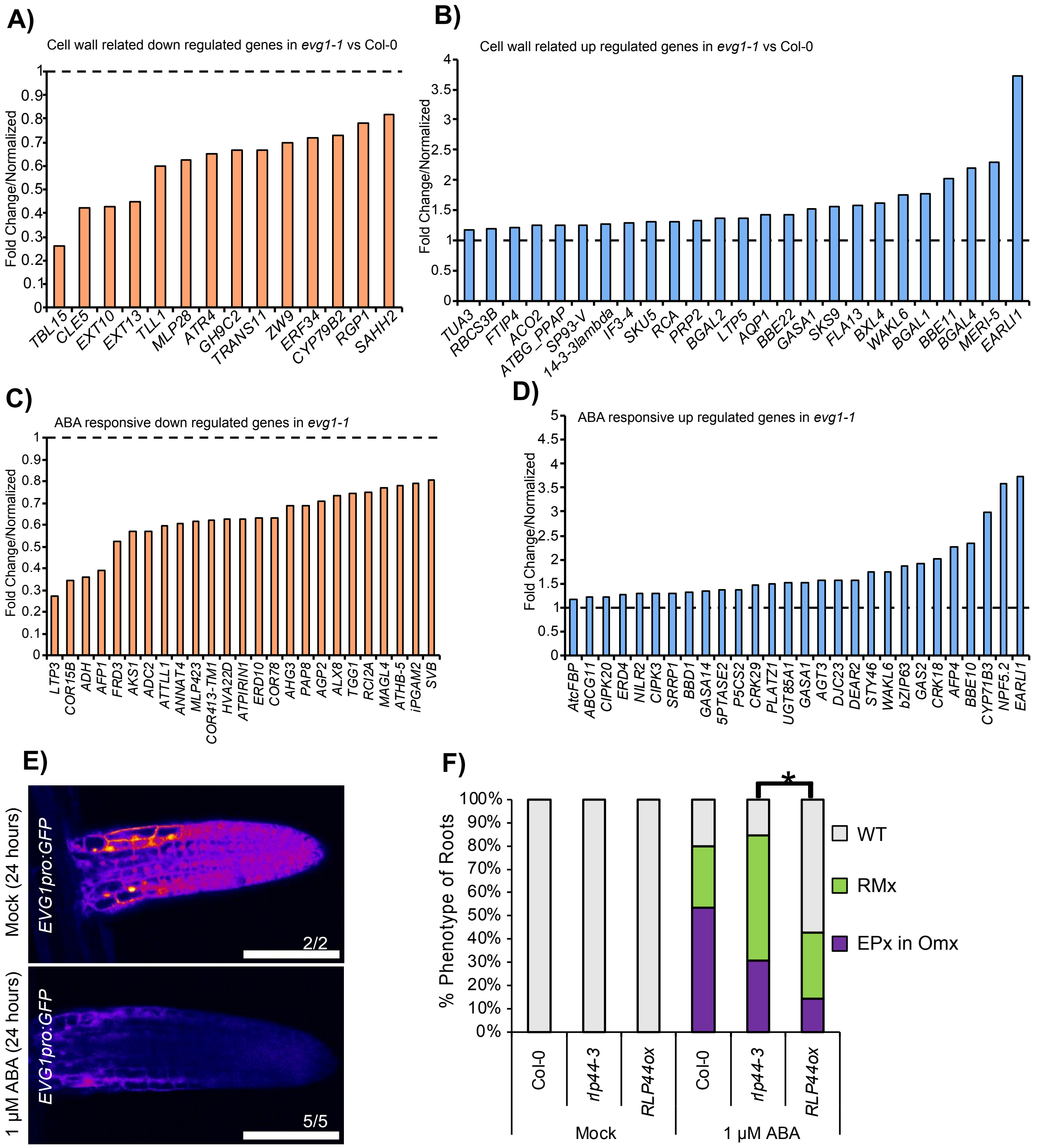


**Supplemental Figure 5. ABA and cell wall effects in *evg1*.** (A-B) Cell wall related genes differentially expressed up or down in *evg1-1*, normalised to wild type Col-0 (C-D) ABA response related genes differentially expressed up or down in *evg1-1*, normalised to wild type Col-0 (E) *EVG1pro:GFP* fluorescence in lateral roots under mock and 1 μM ABA treatments at 24 hours. Scale bars represent 100 μm. Numbers in the picture denotes number of lateral roots observed. (F) Changes in xylem morphology in Col-0 (n=13) *rlp44-3* (n=14) and *RLP44ox* (n=13) roots in mock or with 1 μM ABA treatment. Asterisks indicate significant difference. * $p<0.05$, Fisher's exact test with Benjamini-Hochberg correction.

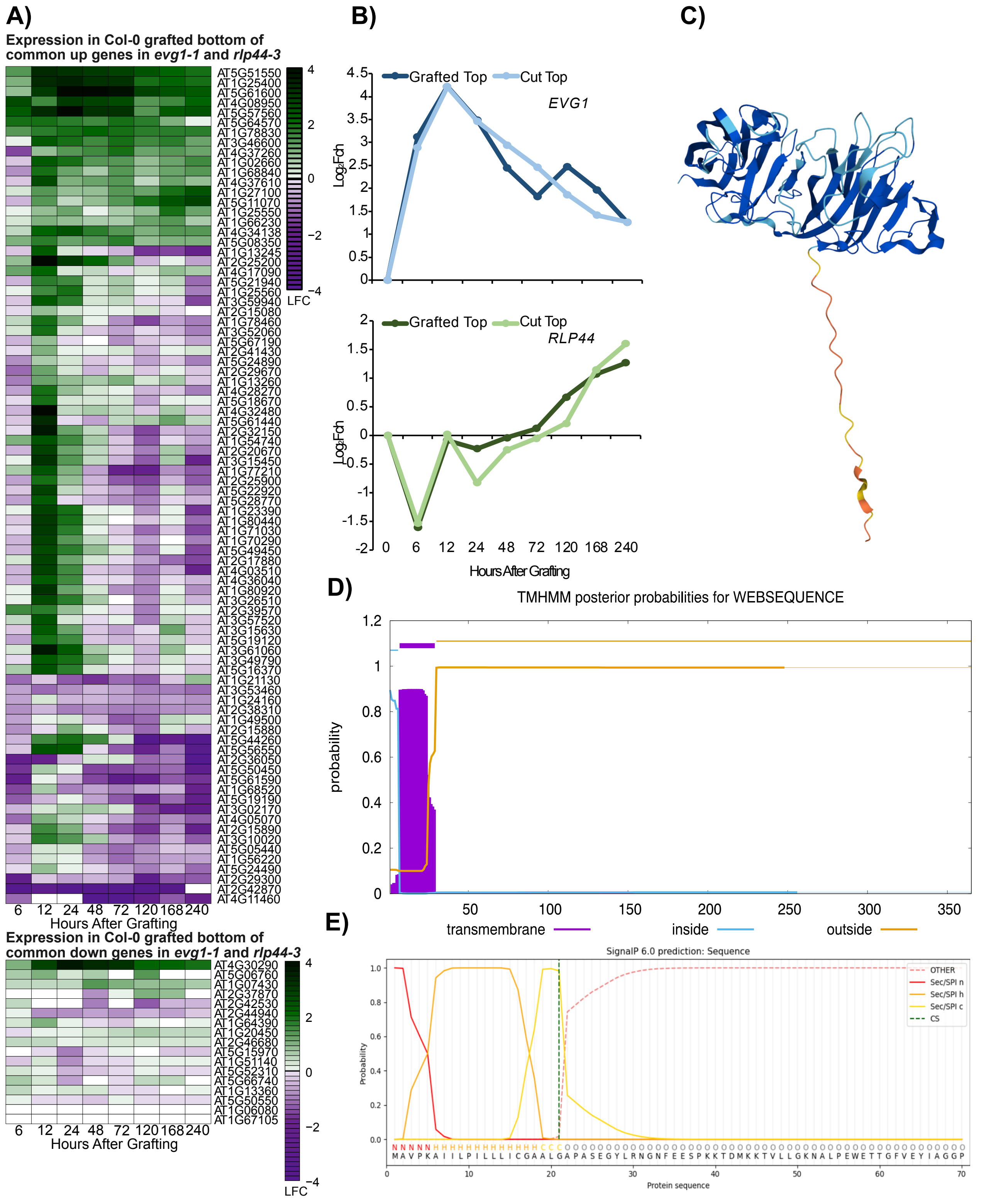


**Supplemental Figure 6. *EVG1* is stress responsive and possesses a signal peptide and a transmembrane domain.** (A) Heat maps with the 84 commonly increased transcripts or 17 commonly decreased transcripts from *evg1-1* and *rlp44-3* showing their expression in wild-type Col-0 grafted bottom tissues during graft formation compared to ungrafted tissues (Melnyk et al., 2018). (B) Comparison of *EVG1* and *RLP44* expression in grafted top and cut top tissues compared to ungrafted during graft formation (Melnyk et al., 2018). (C) Predicted protein structure of EVG1 using AlphaFold 2.0 (Jumper et al., 2021; Varadi et al., 2021). (D) Transmembrane domain prediction using TMHMM (Krogh et al., 2001). (E) Signal sequence prediction using Signal IP6.0 (Teufel et al., 2022).
