## Supplemental Tables 1-4 for "Damage activates *EVG1* to suppress vascular differentiation during regeneration in *Arabidopsis thaliana*"

| **AGI code** | **Name** | **Line** | **Mutant** | **Source** | **Donated by** |
| --- | --- | --- | --- | --- | --- |
| AT3G08030 | *EVG1* | *evg1-1* | SALK_011516C | This Study | NASC |
| AT3G08030 | *EVG1* | *evg1-2* | SALK_119379C | This Study | NASC |
| AT3G08030 | *EVG1* | *EVG1-OE* | - | (Ichikawa et al., 2006) | RIKEN BRC |
| AT3G08030 | *EVG1* | *EVG1pro:GFP* | - | This Study | - |
| AT3G08030 | *EVG1* | *EVG1pro:EVG1-GFP* |  | This Study | - |
| AT3G49750 | *RLP44* | *rlp44-3* | SAIL_596_E12 | (Wolf et al., 2014) | Sebastian Wolf |
| AT3G49750 | *RLP44* | *RLP44ox* | - | (Wolf et al., 2014) |  |
| AT4G32880 | *ATHB8* | *athb8-12* | - | (Prigge et al., 2005) | Annelie Carlsbecker |
| AT5G51550 | *EXL3* | *exl3* | SALK_070475C | This Study | NASC |
| AT3G22600 | *LTPG5* | *ltpg5* | SALK_007462C | (Edstam and Edqvist, 2014) | NASC |
| AT5G10930 | *CIPK5* | *cipk5* | SALK_063555C | (Förster et al., 2019) | NASC |
| AT4G24130 | *SVB5* | *svb5* | SALK_006910C | (Hussain et al., 2021) | NASC |
| AT4G39400 | *BRI1* | *bri1-301* |  | (Li and Nam, 2002) | Sebastian Wolf |
| AT4G18710 | *BIN2* | *bin2-1* |  | (Li et al., 2001) |  |
| AT1G19350 | *BES1* | *bes1-2* | WiscDsLox246D02 | (Lachowiec et al., 2013) | Yuki Kondo |
| AT1G19350 | *BES1* | *bes1-D* |  | (Yin et al., 2002) |  |
| AT1G75080 | *BZR1* | *bzr1-D* |  | (Wang et al., 2002) |  |

Supplemental Table 1: List of lines used and generated in this study

Supplemental Table 2: Primers used for genotyping

| Name | Sequence | Reason |
| --- | --- | --- |
| Evg1-F1 | CAAAGACGACAAAATTCCCAC | Genotyping *evg1-1* |
| Evg1-R1 | CGTGAGTGCGTAGAGAGAACC |  |
| Evg1-F2 | CACTAACCAAGGAAAACTCATTCTC | Genotyping *evg1-2* |
| Evg1-R2 | CAAGCTCTTTAATGGCGACAG |  |
| Exl3-F1 | GCAATCATTAGGCAAAAGCTG | Genotyping *exl3* |
| Exl3-R1 | TCAGCCGTTGGATCTAAACAC |  |
| CIPK5-F1 | GGAAAGTACGAGATGGGAAGG | Genotyping *cipk5* |
| CIPK5-R1 | TGTTCATCTCCTTCGTAACCG |  |
| LTPG5-F1 | TGGGATGCGTACATGTGTATG | Genotyping *ltpg5* |
| LTPG5-R1 | ATGTAGTTGAGACACGGCGAC |  |
| SVB5-F1 | TTGCTTCATCCAAACGTAACC | Genotyping *svb5* |
| SVB5-R1 | AAGGGCTTTGGACTTTACCTG |  |
| RLP44-3-F1 | ATTCCACTCCCAAGTCCAATC | Genotyping *rlp44-3* |
| RLP44-3-R1 | AATGGATGGCATGATTAGGATC |  |
| SALK_LB1 | ATTTTGCCGATTTCGGAAC | LBb1.3 SALK pROK2 |
| SAIL_LB1 | GCCTTTTCAGAAATGGATAAATAGCCTTGCTTCC | LB1 for SAIL lines C/418-451 of pCSA110-pDAP101_T-DNAs |
| pDs-Lox_LB1 | AACGTCCGCAATGTGTTATTAAGTTGTC | WiscDsLox T-DNA LB primer P745 |

Supplemental Table 3: Primers used for expression analysis

| Name | Sequence | Reason |
| --- | --- | --- |
| Evg1_qP-F | GCTCCTGCTTCTGAAGGTTATC | qPCR primers for EVG1 |
| Evg1_qP-R | AACCGGTGGTTTCCCATTCG |  |

Supplemental Table 4: Primers used for Cloning

| Name | Sequence | Cloning module | Reason |
| --- | --- | --- | --- |
| EVG1-proF | AACAGGTCTCAACCTCACCGATGGTGACATTG | Greengate | *EVG1* promoter |
| EVG1-proR | AACAGGTCTCATGTTTGTCTCTGTTGTTCTTCC | Greengate |  |
| RbcsT-F | AACAGGTCTCACTGCAGAGCTTTCGTTCGTATC | Greengate | Pea Rbcs9-E terminator |
| RbcsT-R | ACAAGGTCTCATAGTGTTGTCAATCAATTGGC | Greengate |  |
| GFP-CDS-F | AACAGGTCTCAGGCTATGGTGAGCAAGGGCG | Greengate | GFP coding sequence |
| GFP-CDS-R | ACAAGGTCTCACTGACTACTTGTACAGCTCGTCC | Greengate |  |
